# Endogenous variability in transcription factor concentrations shapes their genome-wide occupancy

**DOI:** 10.1101/2025.09.24.678270

**Authors:** Romane Mizeret, Armelle Tollenaere, Cédric Deluz, David M. Suter

## Abstract

The control of transcription factor (TF) concentrations is crucial for the precise regulation of gene expression and cell fate decisions during development. Yet, TF concentrations can display substantial temporal fluctuations and intercellular variations. How TF levels quantitatively shape genome-wide occupancy patterns remains largely unexplored. Here, we systematically investigate how physiological fluctuations in the concentrations of the pluripotency TFs OCT4, SOX2, and NANOG impact their genomic binding profiles in mouse embryonic stem cells. We uncover distinct concentration-dependent binding behaviors for each TF. NANOG occupancy increased monotonously with its concentration, even though a common set of regions made accessible by OCT4 and SOX2 are highly and uniformly bound at all NANOG concentrations. In contrast, SOX2 occupancy does not increase with its concentrations, while OCT4 displayed an intermediate behavior. Our results challenge the notion that TF binding follows simple mass-action dynamics and reveal a core set of pluripotency regions that are bound by all three TFs at all TF concentrations, revealing how pluripotency can be maintained despite marked fluctuations in core pluripotency TFs.

## Introduction

Transcription factors (TFs) are central regulators of gene expression that act within coordinated networks to maintain or change cell identity. Their binding to chromatin is influenced by both intrinsic factors, such as DNA sequence preference and binding affinity, and extrinsic features like chromatin accessibility, which is regulated by nucleosome positioning and histone modifications ^1–3^. Additional determinants include TF cooperativity ^4–6^, competitive interactions ^7^, and differential affinity for target sequences ^8–10^. TF concentration also plays a key role in determining TF occupancy. During early development of *drosophila*, a gradient of the Bicoid TF specifies developmental patterning by differentially regulating target genes expression in a concentration-dependent manner ^11,12^. TFs are heavily over-represented among haplo-insufficient genes in humans, suggesting the importance of tightly controlling their concentration ^13^. A recent study investigated the concentration-occupancy relationship of SOX9 in the neural crest using a rapid depletion approach. SOX9 binding follows two distinct modes of concentration-dependency. A subset of binding sites decreases its occupancy as SOX9 is progressively depleted (“sensitive regions”), while another group of sites (“buffered regions”) is constantly occupied even at low SOX9 concentrations ^14^. This suggests that different target regions of the genome have evolved mechanisms to confer distinct levels of sensitivity to TF dosage.

The depletion approach used in this study ^14^ is particularly powerful to understand TF dosage effects that are relevant to pathological conditions in which TF expression is abnormally low, such as heterozygous loss-of-function mutations of the *sox9* gene causing campomelic dysplasia ^15^ and Pierre Robin syndrome ^16^. However, even in physiological conditions, TF concentrations can fluctuate substantially over time. In mouse embryonic stem cells (mESCs), a network of TFs plays a central role in maintaining pluripotency. NANOG, OCT4 and SOX2 are core pluripotency TFs that can co-occupy regulatory elements, regulate DNA binding of each other and (for OCT4 and SOX2) control chromatin accessibility of thousands of regions in the genome ^1,2,17–21^. All three TFs exhibit substantial fluctuations of their concentrations on time scales of one to a few cell cycles ^22,23^. These fluctuations were shown to directly impact cell fate decisions of mESCs, with high NANOG levels enhancing the maintenance of the naive pluripotent state ^24^ and high endogenous OCT4 and SOX2 levels facilitating lineage specification ^22^.

To understand how NANOG, OCT4 and SOX2 endogenous fluctuations impact cell fate decisions, we dissected how their physiological fluctuations alter their genomic occupancy, revealing distinct, TF-specific concentration-occupancy relationships.

## Results

### NANOG, OCT4 and SOX2 show distinct concentration-occupancy relationships

We decided to separate cell populations according to physiological variations in their endogenous TF levels, followed by ChIP-seq analysis. This strategy allows to assess concentration-dependent TF binding within the normal TF expression range, thereby avoiding potential artefactual genomic occupancy changes caused by TF knockdown or overexpression. Briefly, after fixation and permeabilization, cells were immunostained for either NANOG, OCT4 or SOX2 and labelled with Hoechst. We then sorted only cells in G1 phase based on Hoechst staining in order to exclude bias from cell cycle phase distribution of the cells sorted for TF levels, which typically double during each cell cycle ^22^. Cells were sorted into four equally-sized populations based on fluorescence intensity, which we used as a proxy for TF concentration (**Methods, Figure 1A**). We then performed calibrated ChIP-seq using drosophila chromatin spike-in (**Methods**) for the corresponding TFs to robustly quantify concentration-dependent changes in specific site occupancy. Relative TF levels were assessed by the average fluorescence intensity of each sorted group. The population with the lowest fluorescence was set as the reference (1X), and fold-changes in TF concentrations were calculated relative to this baseline (**Figure 1B-D**). Fold-change values remained consistent across replicates (**Figure S1A-C**), indicating high reproducibility of the staining and sorting procedures.

**Figure 1.**
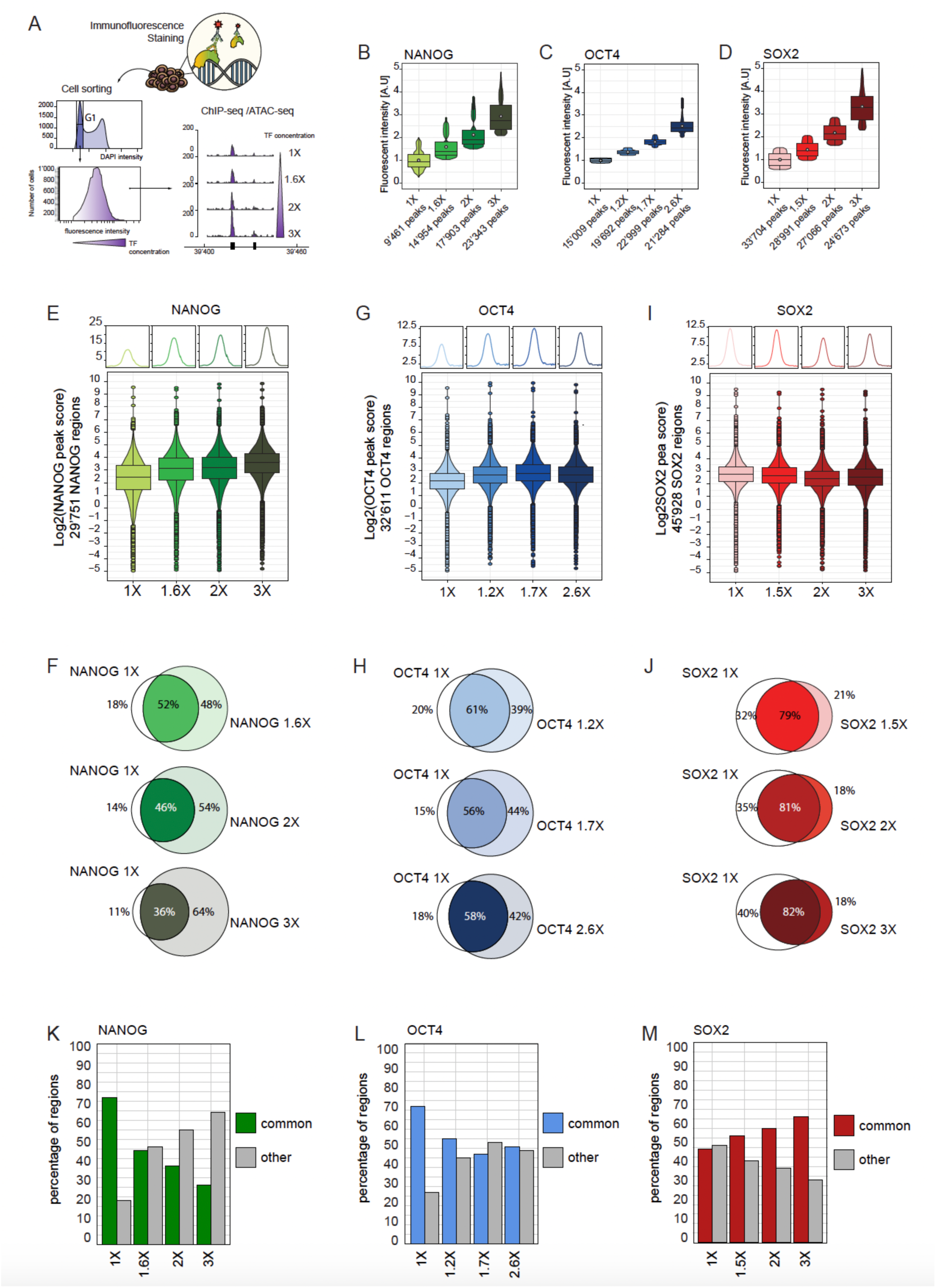
OCT4 and NANOG binding site occupancies scale with their concentrations, while SOX2 binding is concentration independent. (A) Schematic representation of the experimental set up. WT cells were collected, fixed, and immuno-fluorescently stained for OCT4, SOX2 and NANOG then G1 cells were sorted into four distinct concentration binds based on fluorescence intensity (TF concentration). (B-D) Distribution of fluorescence intensity in the sorted populations (n=2) for NANOG (B), OCT4 (C), and SOX2 (D). The baseline average signal was set at 1 (gate 1X), and relative thresholds were calculated for the incrementing gates. The white dots represent the average signal intensity per bin. The edges of the box represent respectively the first and third quartiles (Q1 and Q3), the bar represent the median value and the dots are the outlier values (lower than Q1-1.5xInterquartile-range, higher than Q3+1.5xInterquartile-range) (E, G, I) Representation of OCT4 (E), NANOG (G), and SOX2 (I) binding site occupancies as their concentrations increase. Boxplots show the log2 fold-change in transcription factor (TF) peak scores at their respective target regions. (F, H, J) Overlap of bound regions at different concentrations for NANOG (F), OCT4 (H) and SOX2 (J). (K-M) Fraction of commonly-bound and uniquely-bound regions for NANOG (K), OCT4 (L) and SOX2 (M)

We initially hypothesized that global TF binding site occupancy would scale with their concentration. We found that NANOG occupancy rose monotonously with its concentration both in terms of average peak score and number of bound sites (**Figure 1E-F**). To confirm our results using an approach independent of intracellular antibody staining, we generated a homozygous knock-in cell line in which both alleles of NANOG are fused to YPet (also note that this cell line was also engineered to express mCherry from the endogenous *rex1* locus, but we did not further use this reporter in the present work (**Methods**)). We then sorted cells in four incremental YPet intensity bins (**Figure S1D**) and performed ChIP-seq against NANOG on chromatin extracted from each population. We found a similar concentration-occupancy relationship compared to the one obtained with intracellular antibody staining (**Figure S1E-F**). In contrast to NANOG, OCT4 binding occupancy initially increased concomitantly with its concentration, then reached a plateau, and its number of bound regions dropped slightly at its maximal concentration (**Figure 1G-H**). SOX2 binding site occupancy deviated even more from our expectations, as it slightly decreased with increasing concentrations (**Figure 1I**), and the number of bound regions showed only moderate variation across the four concentration bins (**Figure 1J**). The distribution of TF binding across different types of regions (promoters, genic, intergenic, etc.) only changed marginally as a function of their concentration (**Figure S1G-I**). We next asked whether different bound regions show different TF concentration-dependencies. We identified genomic regions bound at all concentrations regardless of peak score, and calculated their proportion relative to the total bound fraction for each TF. While both NANOG (**Figure 1K**) and OCT4 (**Figure 1L**) displayed an increase in newly occupied regions at increasing concentrations, SOX2 showed the opposite trend (**Figure 1M**). We next looked for enrichment of TF motifs at regions bound by NANOG, OCT4 and SOX2. We observed increasing enrichment for the OCT4:SOX2 and KLF motifs at increasing NANOG concentration, in line with high NANOG concentrations favoring pluripotency maintenance (**Figure S1J**) ^25,26^. Across OCT4-bound sites, the set of enriched motifs remained consistent, with a modest increase in the enrichment of the composite OCT4::SOX2 motif as OCT4 levels increased (**Figure S1K**). In contrast, SOX2-bound regions displayed stable OCT4::SOX2 enrichment profiles and a progressive decrease in KLF motifs with increasing concentrations (**Figure S1L**), in line with high SOX2 levels favoring cell differentiation ^22^.

### Chromatin accessibility and other TFs shape concentration-dependent NANOG occupancy

We next investigated whether TF concentration-occupancy varies as a function of chromatin accessibility using previously published ATAC-seq data ^2^. NANOG showed a shift to less accessible sites with increasing concentrations (**Figure 2A**). In contrast, both OCT4 (**Figure 2B**) and SOX2-bound regions (**Figure 2C**) displayed relatively consistent levels of accessibility across all concentrations.

**Figure 2.**
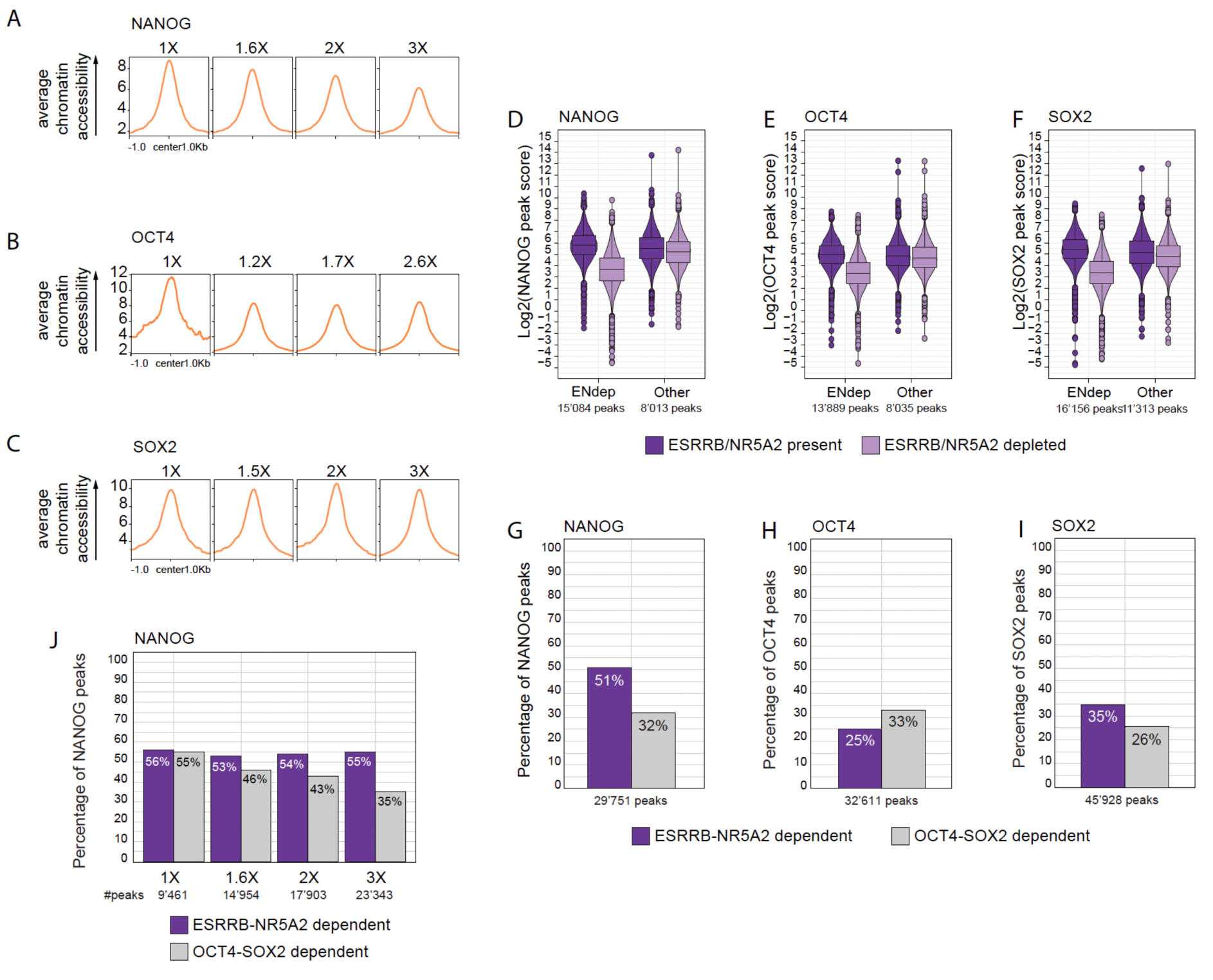
NANOG binds less accessible sites as its concentration increases, while OCT4 and SOX2 target equally accessible regions independently of their levels. (A-C) Average chromatin accessibility (RPKM normalized) at regions bound by NANOG (A), OCT4 (C), and SOX2 (C), centered on their binding sites, across their increasing concentrations. (D-F) Distribution of peak scores for NANOG (D), OCT4 (E), and SOX2 (F) at ESRRB-NR5A2-dependent or independent regions, in the presence (dark purple) or absence (light purple) of ESRRB and NR5A2. (G-I) Proportion of OCT4-SOX2-dependent and ESRRB-NR5A2-dependent regions within the target sites of NANOG (G), OCT4 (H), and SOX2 (I). (J) Proportion of OCT4-SOX2-dependent and ESRRB-NR5A2-dependent regions in NANOG-bound sites as its concentration increases. (ATAC-seq data from Placzek et al.,2025, ESSRB-NR5A2 data from Festuccia et al., 2021, OCT4-SOX2 data from Friman et al., 2019).

We next asked to which extent NANOG depends on other TFs for its binding, when NANOG is expressed at different concentrations. It is known that OCT4, SOX2, ESRRB and NR5A2 can help NANOG binding ^2,27^. Specific regions of the genome require continuous binding of either OCT4, SOX2, or both factors to remain accessible ^28^, and we are referring to these regions as OCT4-SOX2-dependent. ESRRB and NR5A2 also regulate NANOG (and OCT4 and SOX2) binding at co-bound targets ^29^, and were suggested to maintain chromatin accessibility at their target sites ^29,30^. Using data from ^29^, we identified regions for which OCT4, SOX2 and NANOG binding drops by two-fold or more upon ESRRB and NR5A2 co-depletion (**Figure 2D-F**), and defined these as ESRRB-NR5A2 dependent. We measured the proportion of OCT4-SOX2 and ESSRB-NR5A2 dependent regions among bound sites. 51% of NANOG-bound sites depended on ESRRB-NR5A2 (**Figure 2G**), while OCT4 and SOX2 relied on ESRRB-NR5A2 for their binding at only 25% (**Figure 2H**) and 35% (**Figure 2I**) of their regions, respectively. We next asked how the relative dependency of NANOG for OCT4-SOX2 vs ESRRB-NR5A2 varied as a function of NANOG levels. We found that as NANOG concentration increased, the fraction of occupied sites that were OCT4-SOX2 dependent decreased, while it remained constant for ESRRB-NR5A2 dependent regions (**Figure 2J**). Therefore, as NANOG concentration increases, it becomes less reliant on chromatin accessibility and OCT4-SOX2 for its binding, while retaining the same reliance on ESRRB-NR5A2 co-binding.

We next classified NANOG-bound regions as a function of their concentration dependency. Inspired by previous work dissecting SOX9 concentration-occupancy relationship ^14^, we defined buffered regions as those that were insensitive to NANOG concentration for their occupancy, and sensitive regions as those that increased occupancy with concentration. This approach identified 5,894 buffered regions and 23,857 sensitive regions (**Figure 3A and Figure S2A**). NANOG bound more strongly to buffered regions than to sensitive regions at all concentrations (**Figure 3A**). Buffered regions were also more accessible than sensitive regions (**Figure 3B**) and represented a large fraction of NANOG-bound regions at low concentration, but this fraction progressively declined as NANOG levels increased (**Figure S2B**). Buffered and sensitive regions were similarly enriched at promoters and enhancers (**Figure S2C**). TF motifs that were enriched in NANOG-sensitive and NANOG-buffered regions were highly similar (**Figure S2D-E**). The motif strength for NANOG, OCT4::SOX2, SOX2, ESRRB and NR5A2 showed no marked difference, while the OCT4 motif was weaker in the sensitive regions (**Figure S2F**).

**Figure 3.**
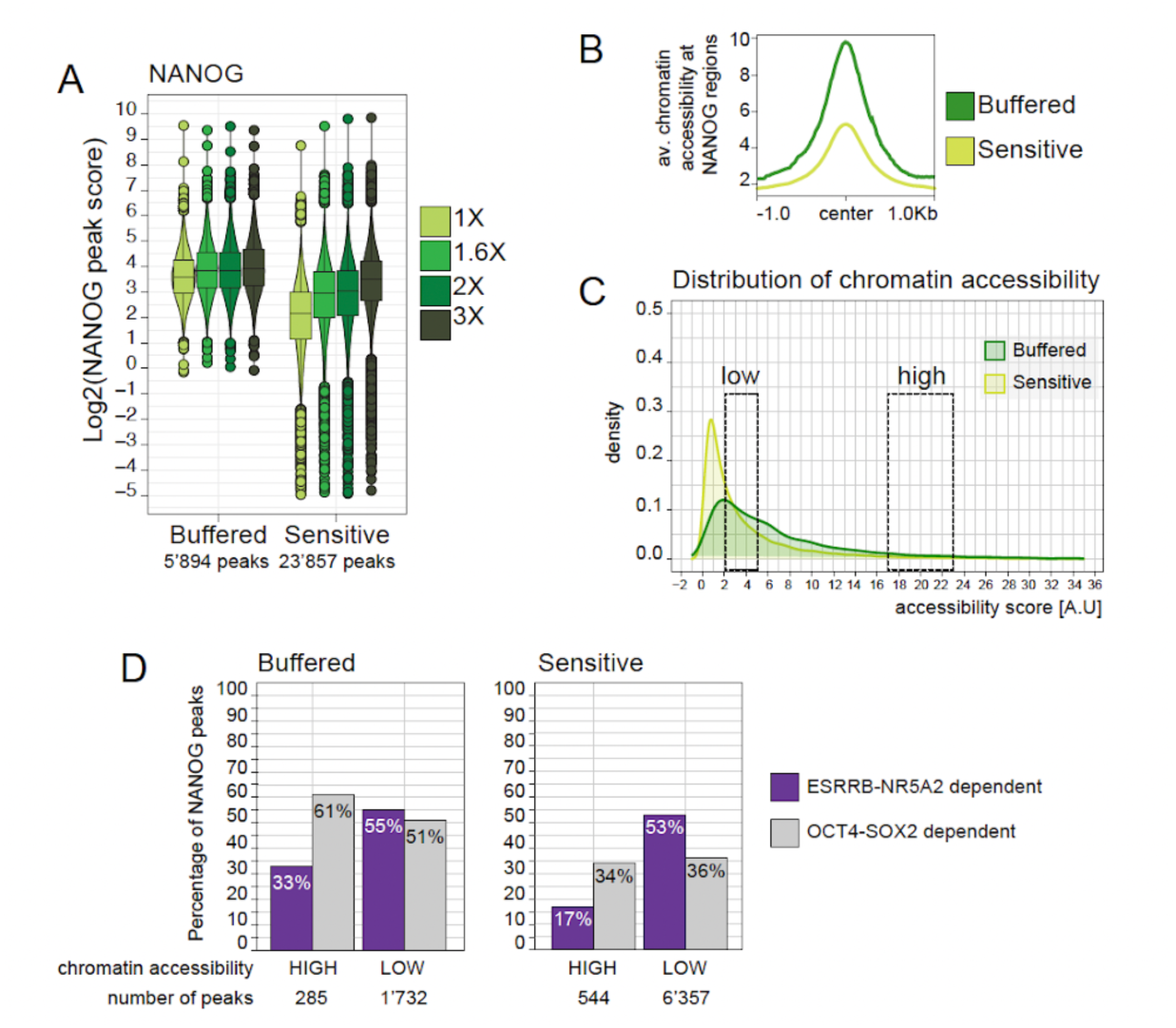
NANOG binds stably at highly accessible, OCT4-SOX2-enriched regions, while ESRRB-NR5A2 assists NANOG binding at less accessible regions. (A) Distribution of NANOG peak scores at NANOG-buffered and NANOG-sensitive regions at increasing concentrations. Boxplots as in Figure 1. (B) Average chromatin accessibility (RPKM normalized) profile of NANOG-buffered (dark green) and NANOG-sensitive (light green) regions centered around NANOG binding sites. (C) Density plot representing chromatin accessibility distribution for NANOG-buffered (dark green) and NANOG-sensitive (light green) regions. Black boxes: selected groups of high and low accessibility regions. (D) Proportions of ESRRB-NR5A2-dependent (purple) and OCT4-SOX2-dependent (gray) regions in NANOG-buffered and NANOG-sensitive regions at high and low accessibility (ATAC-seq data from Placzek et al.,2025, ESSRB-NR5A2 data from Festuccia et al., 2021, OCT4-SOX2 data from Friman et al., 2019).

To explore what distinguishes buffered from sensitive regions beyond accessibility, we asked whether binding of ESRRB/NR5A2 or OCT4/SOX2 within the same regions than NANOG could make NANOG binding insensitive to its concentration at buffered regions. To this end, we separated NANOG-bound regions into high and low accessibility groups using previously published ATAC-seq data ^2^ (**Figure 3C**). We selected groups with comparable median and mean accessibility values across groups (**Figure S2G**). We then quantified the proportion of ESRRB-NR5A2 dependent and OCT4-SOX2 dependent regions within these subgroups. ESRRB and NR5A2 dependent regions were enriched at NANOG-bound sites with low accessibility, represented at 55% and 53% of lowly accessible NANOG buffered regions and sensitive regions, respectively, but only 33% and 17% of highly accessible buffered and sensitive regions, respectively (**Figure 3D**). In contrast, OCT4-SOX2 dependent regions represented 61% and 51% of NANOG buffered regions with high and low accessibility, respectively, but only 34% and 36% of NANOG sensitive regions, respectively (**Figure 3D**). Together, these results suggest that ESRRB and NR5A2 support NANOG binding at less accessible regions, while OCT4 and SOX2 co-binding buffers a subset of accessible regions against changes in NANOG concentration ^22^.

### Concentration-dependent and SOX2-dependent binding of OCT4

To dissect the concentration-dependency of OCT4 binding, we first attempted to group regions as buffered or sensitive based on OCT4 peak scores as we did for NANOG. However, both regions classified as OCT4-buffered and OCT4-sensitive regions exhibited similar occupancy profiles across concentrations, with a peak at intermediate levels followed by a decrease in occupancy at higher OCT4 levels. Although concentration-dependent changes in OCT4 occupancy were more pronounced in sensitive regions, the overall similarity of these profiles suggests that this classification does not reflect biologically significant, distinct OCT4 binding behaviors (**Figure S3A**).

To identify regions where OCT4 binding responds in a concentration-dependent manner, we applied unsupervised clustering to OCT4-bound regions and defined three clusters (**Methods**, **Figure S3B-C**). In cluster O1, OCT4 binding peaked at 1.2X and decreased at higher concentrations. In cluster O2, binding increased up to 1.7X, while in cluster O3, binding progressively increased across the full concentration range without plateauing (**Figure 4A**). We also found that OCT4 only slightly redistributed to promoter regions at increasing concentrations (**Figure S3D**). These patterns suggest that OCT4 binds distinct subsets of regions depending on its concentration and partially redistributes as its concentration varies. We next examined whether these clusters differ in chromatin accessibility or in the nature of the bound TF motifs. Despite reaching saturation at lower concentrations, O1 regions were on average less accessible than those in clusters O2 and O3 (**Figure 4B**). OCT4-SOX2 dependent regions accounted for 40% of cluster O1, compared to only 30% and 26% of clusters O2 and O3, respectively (**Figure 4C**). O1 regions were also more enriched for the OCT4::SOX2 motif (**Figure S3E**). We next assessed OCT4 binding upon SOX2 depletion using previous datasets that we had acquired in the presence or absence of SOX2 in the 2TS22C cell line ^31,32^. We found that OCT4 binding decreased more markedly at O1 regions upon SOX2 depletion than in the other clusters (**Figure 4D**), confirming that SOX2 contributes specifically to OCT4 binding at less accessible, co-bound targets and helps maximizing OCT4 binding in O1 regions at low Oct4 concentrations.

**Figure 4.**
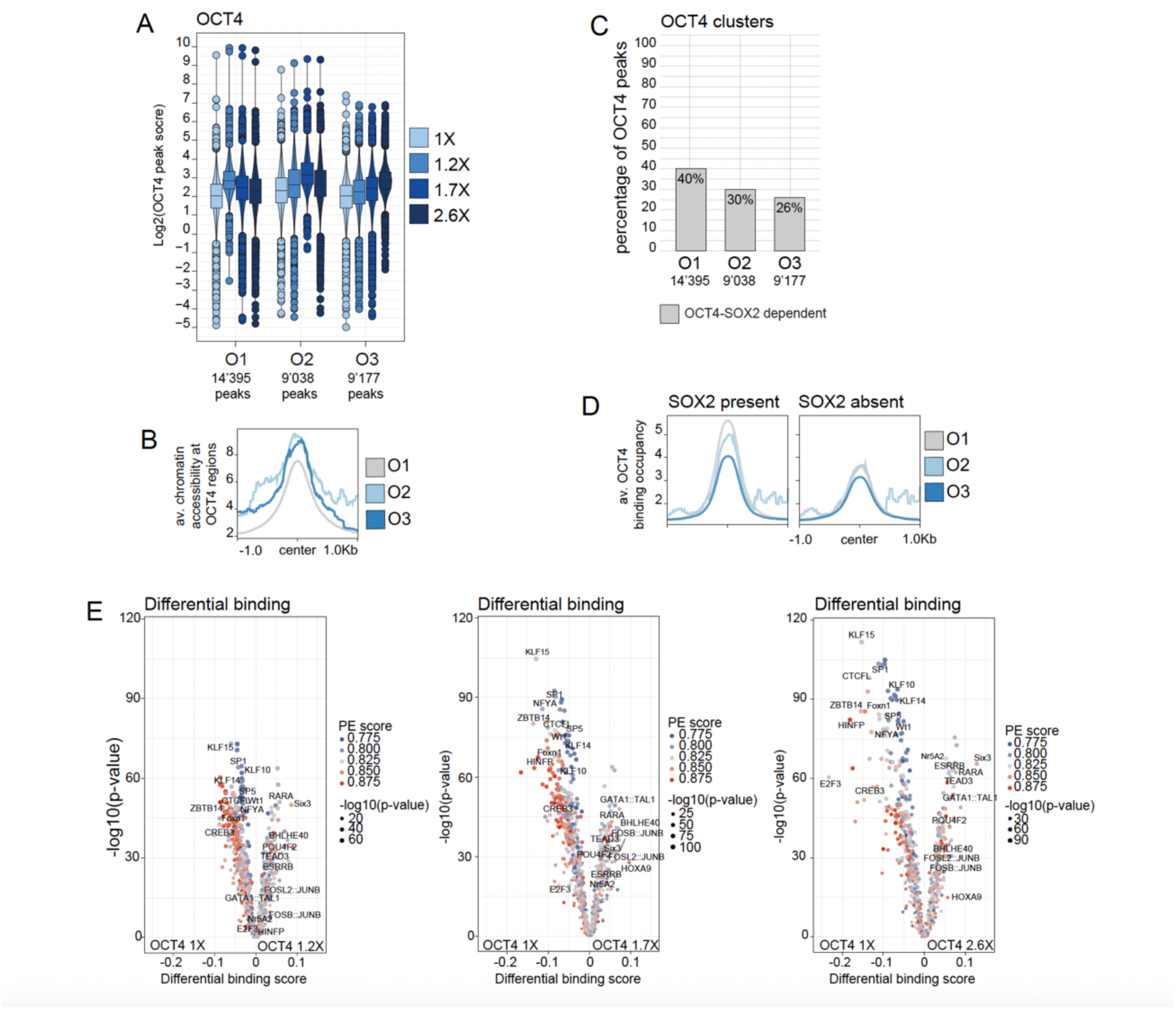
OCT4 binding to less accessible regions is more dependent on SOX2 than in more accessible regions. (A) Boxplot displaying the distributions of OCT4 peak scores across OCT4-bound regions, grouped into cluster O1 (14,395 peaks), O2 (9,038 peaks), and O3 (9,177 peaks). Boxplots as in Figure 1 (B) Average chromatin accessibility (RPKM normalized) of OCT4-bound regions at each cluster, centered around OCT4 binding sites. (C) Number of OCT4 regions that are dependent on ESRRB-NR5A2 and OCT4-SOX2 within each OCT4 cluster. (D) Profiles depicting OCT4 peak scores at OCT4-bound regions per clusters when SOX2 is present (right) or depleted (left) (Placzek et al., 2025, Cell Rep.). (E) TF footprints enrichment at OCT4-bound regions with increasing OCT4 concentrations compared to baseline (OCT4 1X), using TOBIAS (Bentsen, M et al. 2020). TFs are assigned a promoter-enhancer score (PE score) based on whether their motifs occur at promoters (red dots) or enhancers (blue dots) (ATAC-seq data from Placzek et al.,2025, ESSRB-NR5A2 data from Festuccia et al., 2021, OCT4-SOX2 data from Friman et al., 2019).

We had previously reported that cells with high endogenous OCT4 concentrations display increased chromatin accessibility at differentiation loci ^22^. This prompted us to ask whether different endogenous concentrations of OCT4 would result in differential binding of other TFs, in particular those involved in differentiation. We thus sorted cells based on OCT4 levels, matching normalized OCT4 concentrations of 1X, 1.5X, 2X and 2.6X. We then performed ATAC-seq on fixed cells isolated from each bin, and performed TF footprinting analysis using the TOBIAS pipeline ^33^. We then compared the TF footprinting scores of OCT4 1.5X, 2X and 2.6X with the OCT4 1X bin. Footprints with differential scores were consistent among these and the differences increased with larger differences in OCT4 concentrations (**Figure 4E**). With increasing OCT4 concentrations, footprints for TFs of the KLF family, NFY and SP1 decreased, while footprints of TFs linked to differentiation such as Jun/Fos, GATA1, Retinoic acid receptor and TEADs increased (**Figure 4E**). Therefore, even though OCT4 binding plateaus at intermediate concentrations, its impact on binding of other TFs monotonously increases with its concentration. These changes in TF binding are also in line with our previous work showing that higher OCT4 concentrations prime mESCs for differentiation ^22^.

### Concentration-dependent SOX2 redistribution away from OCT4::SOX2 motifs

Unlike OCT4 or NANOG, SOX2 binding occupancy did not scale with its concentration and was mostly bound to a common set of regions across all SOX2 levels (**Figure 1E**). To further investigate how SOX2 concentration regulates its binding at target sites, we clustered SOX2-bound regions based on Z-score normalized peak scores. Gap statistics identified a plateau around three clusters (**Figure S4A-B**), which we retained for further analysis. In cluster S1, SOX2 binding peaked at low concentrations while in cluster S3, binding peaked at the highest concentration (**Figure 5A).** Chromatin accessibility was highest in cluster S1 and lowest in cluster S3 (**Figure 5B**). Cluster S2 displayed an intermediate behavior for these parameters (**Figure 5A-B**). OCT4-SOX2-dependent regions were most enriched in clusters S1 and S2 (27% and 29%, respectively) and least in cluster S3 (19%) (**Figure 5C**). All SOX2 clusters were similarly enriched for previously described ESRRB-NR5A2-dependent regions (**Figure S4C**), suggesting that ESRRB and NR5A2 do not regulate SOX2 binding in a concentration-dependent manner.

**Figure 5.**
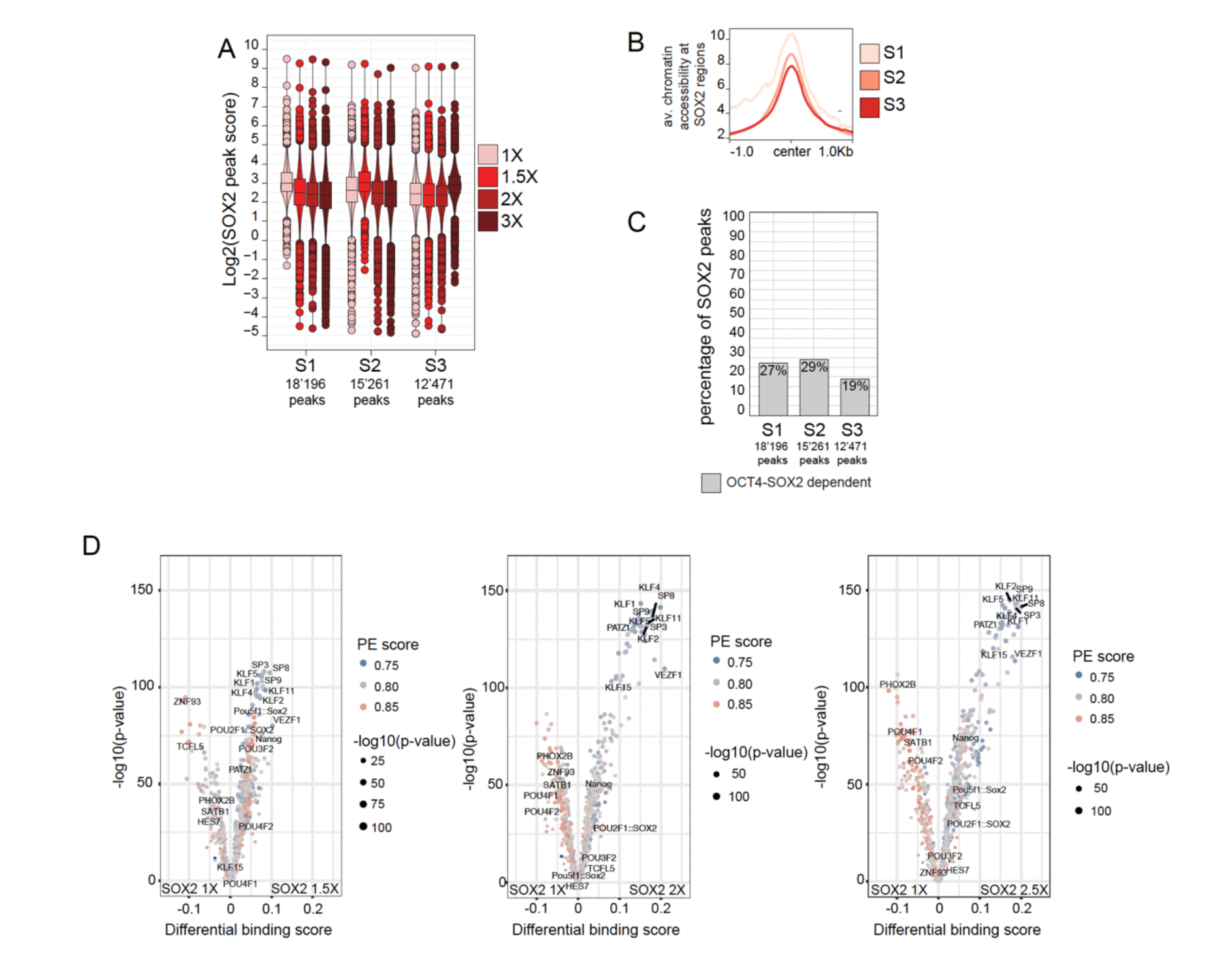
At higher concentration, SOX2 is redistributed to closed chromatin regions enriched at promoters and SOX2 promotes TFs binding to promoters as its concentration increases. (A) Distributions of SOX2 peak scores across SOX2-bound regions, grouped into clusters S1 (15,261 peaks), S2 (12,471 peaks), and S3 (18,196 peaks). (B) Average chromatin accessibility of SOX2-bound regions (45’928 peaks) grouped by cluster, centered on SOX2 motifs, RPKM normalized, data from Placzek et al., 2025. (C) Number of SOX2-bound regions dependent on ESRRB and NR5A2 or OCT4 and SOX2, within each SOX2 cluster. ESRRB-NR5A2 data from Festuccia et al., 2021, OCT4-SOX2 data from Friman et al., 2019. (D) TF footprints enrichment at SOX2-bound regions with increasing SOX2 concentrations compared to baseline (SOX2 1X), using TOBIAS (Bentsen et al., 2020). TFs are assigned a promoter-enhancer score (PE score) based on whether their motifs occur at promoters (red dots) or enhancers (blue dots).

We next performed motif enrichment analysis on the different clusters, which revealed that regions in S1 were most enriched for the OCT4::SOX2 motifs and motifs of pluripotency-associated TFs (e.g., KLF2, KLF4, KLF5), while regions in cluster S2 and S3 were more enriched for motifs of TFs involved in differentiation (e.g., SOX4, SOX6, SOX9, SOX10), (**Figure S4D**). We next performed TOBIAS footprinting analysis on SOX2-bound regions using ATAC-seq data from SOX2-sorted cells, as described above for OCT4. We compared the footprinting scores of SOX2 1.5X, 2X and 3X with the SOX2 1X bin. Sorting was carried out following the same protocol and gating strategy used for ChIP-seq. Similar to OCT4, the directionality of footprint changes was consistent among these comparisons, and the differences increased with larger differences in SOX2 concentrations (**Figure 5D**). TF footprinting revealed a concentration-dependent increase in promoter-associated TF binding, particularly for KLF family members, and a decrease in enhancer-associated TF binding (**Figure 5D**). Taken together, our data supports a model in which a larger fraction of SOX2 molecules localize away from the canonical OCT4::SOX2 motif at moderate-to-high SOX2 concentrations. However, high SOX2 concentrations also result in increased TF binding of some TFs that bind mostly to promoters, such as KLF factors. These results are in agreement with the modest impact of high endogenous SOX2 concentrations on fostering neuroectodermal commitment ^21,22^.

### OCT4 and NANOG binding occupancy decreases at high SOX2 concentrations

We next asked how NANOG and OCT4 genomic occupancy varies across SOX2 concentrations. We first performed co-staining of NANOG, OCT4 and SOX2 followed by flow cytometry analysis (**Methods**) to determine how the concentrations of these TFs co-vary. While OCT4 and SOX2 concentrations were mildly positively-correlated, NANOG and SOX2 showed a stronger positive correlation (**Figure 6A**). Based on the OCT4 and NANOG concentration-occupancy relationships we determined, we thus expected that OCT4 occupancy would be modestly correlated to SOX2 concentration, why NANOG occupancy would scale with SOX2 concentration. We then sorted cells according to SOX2 concentrations as before and performed ChIP-seq against OCT4 or NANOG in the four SOX2 concentration bins (**Figure 6B**). Surprisingly, we found that both NANOG and OCT4 binding decreased at increasing SOX2 levels (**Figure 6C-D**). Chromatin accessibility at the bound regions was similar for NANOG at all SOX2 levels (**Figure 6E**), however OCT4 remained bound only to highly accessible regions as SOX2 levels increased (**Figure 6F**). We also found that the motif quality for NANOG, OCT4::SOX2 and SOX2 tended to increase at the bound sites for higher SOX2 concentrations (**Figure 6G-H**). This suggests that NANOG and OCT4 remain bound only to high-affinity, accessible sites when SOX2 concentration is high.

**Figure 6.**
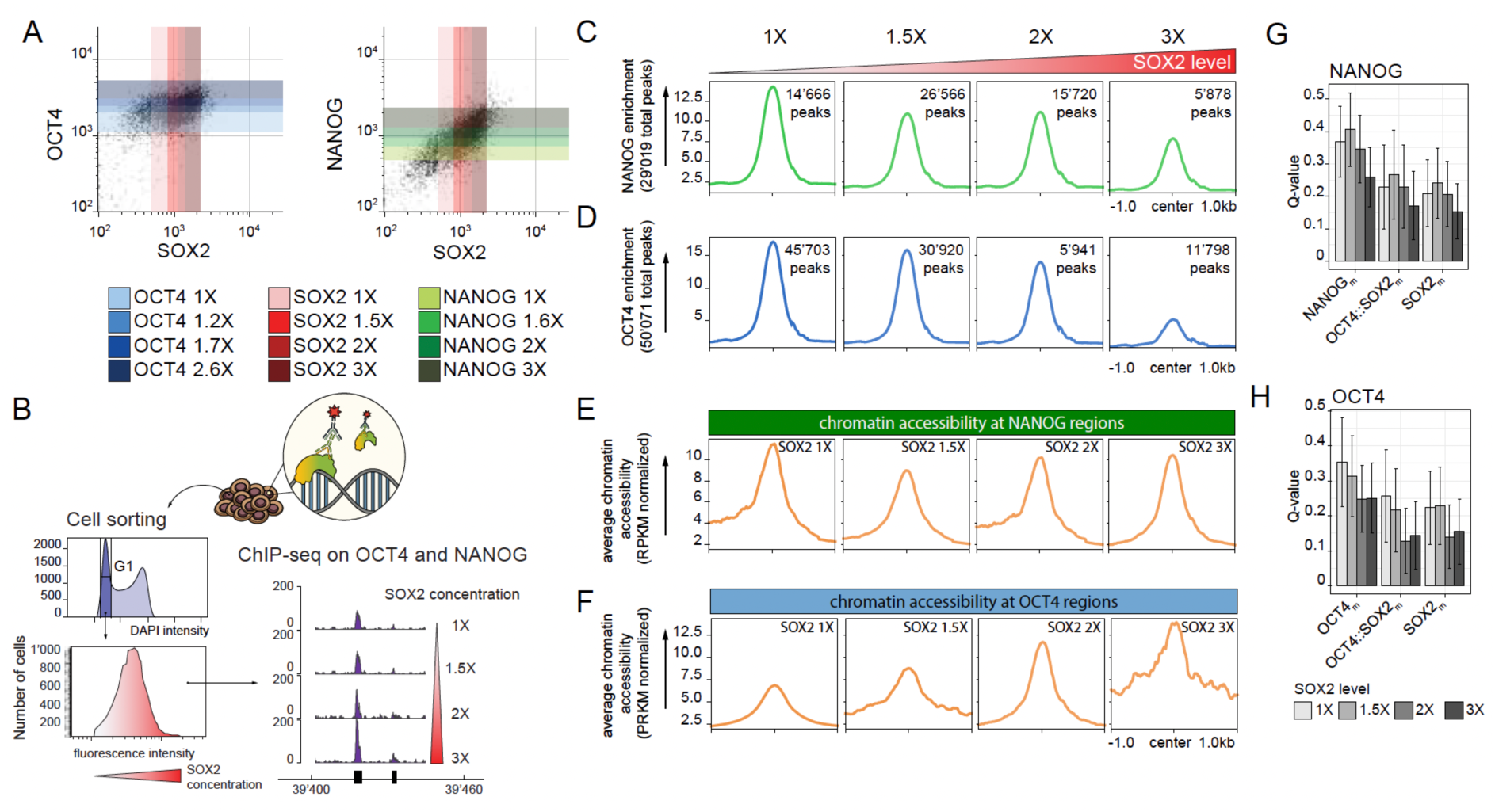
OCT4 and NANOG binding occupancies decrease as SOX2 concentration increases. (A) Correlation between OCT4 vs. SOX2 levels (right) and NANOG vs. SOX2 levels (left) in G1-phase WT mES cells. The color bars represent increasing TF concentrations, mirroring the sorting gates used for TF-concentration-sorted cells (1 replicate). (B) Experimental setup. WT CGR8 cells were fixed, permeabilized, and stained for SOX2. The cells were sorted into four bins based on increasing SOX2 concentration, and OCT4 and NANOG were ChIP-sequenced on the extracted chromatin (1 replicate). (C, D) Binding occupancies of NANOG (C) and OCT4 (D) as a function of SOX2 levels. Enrichment profiles represent the average normalized ChIP-seq signal across all detected binding sites, centered on peak summits (±1kb). Numbers indicated on the top left of each profile represent the number of peaks bound by the TF for a given SOX2 concentration (E, F) Average chromatin accessibility of NANOG-(E) and OCT4-bound (F) regions as a function of SOX2 concentrations. (G-H) Quality scores (Q-values) of OCT4, SOX2, OCT4::SOX2, and NANOG motifs at NANOG- (G) and OCT4-bound (H) regions at varying SOX2 concentrations, as determined using FIMO (Find Individual Motif Occurrences) analysis. A lower Q-value indicates higher confidence in motif quality and a lower false discovery rate (FDR). (ATAC-seq data from Placzek et al., 2025).

### Regions key for pluripotency maintenance are occupied by NANOG, OCT4 and SOX2 at all concentrations

Finally, we asked whether gene regulatory regions that are central to pluripotency maintenance are more likely to be always bound irrespective of TF concentrations. We thus selected regions that are bound by all three TFs i) at all concentrations of all TFs (AC regions) ii) only at some concentrations of the three TFs, or concentration-specific (CS regions). This resulted in 4908 AC and 9881 CS regions that we then further analyzed. For all three TFs and independently of their concentrations, the binding scores were higher in AC regions compared to CS regions (**Figure S5A-C**). AC regions were characterized by a higher enrichment in distal gene regulatory regions (**Figure 7A**), higher occurrences and better motif quality for OCT4::SOX2 and NANOG motifs, and were more enriched in GO terms for stem cell maintenance and PluriNetwork (**Figure 7B-D**). We next assigned AC and CS regions matching gene regulatory sequences to promoters and enhancers of specific genes (**Methods**). There were nearly the same number of regulatory sequences of known regulators of pluripotency enriched in the AC regions (19/29) as compared to the CS regions (20/29), despite the fact that AC regions were linked to about half as many genes as CS regions (2618 vs 4567 genes) (**Figure 7E and Figure S5D**). This suggests that pluripotency network regulatory regions have evolved to be bound by their key regulators independently of TF concentrations.

**Figure 7.**
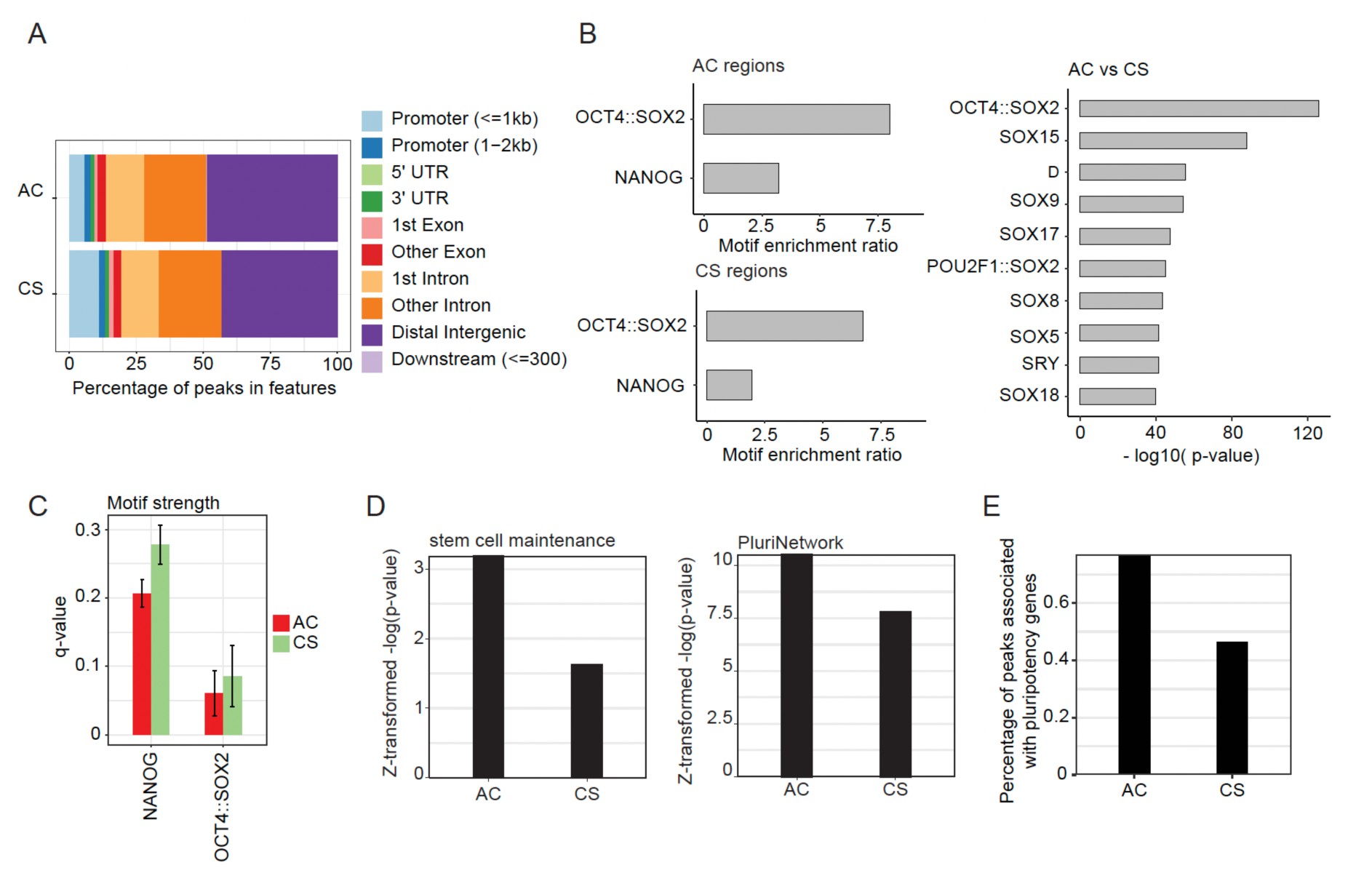
The regions bound by OSN at all concentrations are associated with pluripotency maintenance. (A) Percentage of peaks in AC and CS falling in different genomic features. (B) Left: Enrichment ratio of OCT4::Sox2 and Nanog motifs (determined by SEA) in AC and CS peaks. Right: Comparative motif enrichment analysis of AC versus CS regions. (C) OCT4::Sox2 and Nanog motif strength (resemblance to consensus motif) determined by FIMO in AC and CS regions. A lower q-value indicates higher confidence in motif quality and a lower false discovery rate (FDR) (D) Occurrence of the GO terms stem cell maintenance and PluriNetwork pathway in genes associated with AC and CS peaks. (E) Percentage of peaks predicted to fall into regulatory regions of pluripotency-related genes. Peaks were assigned to the closest gene.

## Discussion

Our study reveals that NANOG, OCT4, and SOX2 each modulate their binding landscape as a function of their endogenous concentration with distinct behaviors. This indicates that TF dosage effects cannot be universally explained by simple mass-action principles even within a physiological TF concentration range. The notion that TF levels modulate target engagement has been well established in the context of *Drosophila* ^12,34^ and *C.elegans* ^35^ development. In this context, higher TF concentrations broaden the binding landscape to include lower-affinity or less accessible sites, whereas reduced concentrations restrict occupancy to highly accessible regions that are also bound by co-factors. Our data indicate that this principle holds true for NANOG, which exhibited a monotonic increase in both peak intensity and number of bound sites with rising concentration. Yet even for NANOG, a large subset of “buffered” regions were highly bound in a concentration-independent manner, similarly to what has been observed for SOX9 concentration-insensitive “buffered” sites described in human cranial neural crest cells ^14^. Buffered NANOG sites were typically accessible and enriched for OCT4 and SOX2 co-binding, pointing to cooperative stabilization as a mechanism that insulates these loci from endogenous NANOG fluctuations. While NANOG binds low-accessibility, pluripotency-associated sites with ESRRB and NR5A2 at high levels, occupancy at these sites diminishes at low NANOG concentrations. This is in line with the lower resistance to differentiation of NANOG-low cells ^36^. The behavior of OCT4 diverged from NANOG with an increase in occupancy at moderate concentrations followed by a plateau accompanied by redistribution among targets. SOX2 exhibited a surprising lack of concentration-dependent occupancy scaling, yet at higher concentrations it shifted towards less accessible chromatin regions. This redistribution occurred away from its canonical OCT4::SOX2 motif and coincided with decreased NANOG and OCT4 occupancy at shared sites, despite unchanged chromatin accessibility. These observations suggest a competitive displacement mechanism, where elevated SOX2 levels redirects its binding away from pluripotency-associated targets and loses its stabilizing effect on NANOG and OCT4 binding at co-bound regions. This change in the SOX2 binding landscape is consistent with previous work linking high SOX2 expression to facilitation of neuroectodermal differentiation ^22^.

Taken together, these observations place pluripotency TF dosage regulation within a broader conceptual framework that has emerged from developmental systems biology. TF binding responses to concentration are highly site- and context-specific, shaped by intrinsic binding affinity, local chromatin accessibility, and co-binding with other TFs. Our results indicate that pluripotency regulatory elements marked by the OCT4::SOX2 heterodimeric motifs emerge as the top motif at all TF concentrations. Therefore, these genomic regions have evolved the ability to maintain TF binding despite marked fluctuations in central pluripotency TFs. Taken together, these results explain how mESCs can maintain pluripotency despite large fluctuations in the levels of core pluripotency TFs.

## Material and Methods

### Cell culture

CGR8 mouse embryonic stem (ES) cells (Sigma, Cat#07032901_1LV) were routinely cultured on 0.1% gelatin-coated (Sigma, G9391) 10 cm Petri dishes. Cultures were maintained at 37°C with 5% CO₂ in N2B27 medium, a 1:1 mixture of Neurobasal Medium (Gibco, 21103049) and DMEM/F-12 (Gibco, 21331020), supplemented with B-27 (Gibco, 17504044), N-2 (Gibco, 17502048), 2 mM L-glutamine (Gibco, 25030-024), 1% penicillin-streptomycin (BioConcept, 4-01F00-H), and 100 μM 2-mercaptoethanol (Sigma, 63689). The medium was further supplemented with 3 μM CHIR99021 (Merck, 361559), 1 μM PD184352 (Sigma, PZ0181), and in-house produced leukemia inhibitory factor (LIF).

Cells were split every 2-3 days based on confluency. To detach cells, Accutase (Brunschwig, ICTAT104) was added to the culture dish, and cells were collected by centrifugation at 1000 x g for 4 minutes at room temperature. Cells were washed in buffer medium before being reseeded, following a previously published, detailed protocol ^37^.

For immunofluorescence imaging, cells were plated on Laminin-coated surfaces (1:10 dilution of Biolaminin LN511-0202, BioLamina in PBS). Prior to imaging, the medium was replaced with phenol-free N2B27 medium (same composition as N2B27) supplemented with 3 μM CHIR99021, 1 μM PD184352, and in-house produced LIF.

### Generating the YPet-Nanog homozygous cell line

The YPet-Nanog cell line was derived from a homozygous CGR8-derived cell line carrying a P2A-mCherry reporter downstream of the naive pluripotency marker *Rex1* (*Zfp42*) coding sequence (**Figure S6A**). Note that we did not use the mCherry reporter in the context of this study, but we still describe the generation of this cell line here as it has not been described in a publication so far. To introduce this reporter, guide RNAs (gRNAs) targeting the REX1 C-terminal were designed using CRISPOR^154^ and cloned into the pX335-gRNA vector (pX335-U6-Chimeric_BB-CBh-hSpCas9n(D10A), a gift from Feng Zhang (Addgene plasmid #42335) by digestion with BbsI, generating the pX335-cas9D10A-Rex1guide4, pX335-cas9D10A-Rex1guide53.

A donor cassette (pSUT1-REX1) containing the P2A–mCherry sequence, fused in frame to a SV40 nuclear localization signal and a PEST degron, flanked by 500 bp REX1 homology arms in which the endogenous stop codon was removed, was ordered from GeneArt. For transfection, 4 million WT GCR8 cells were plated on 0.1% gelatin-coated Petri dishes in 10mL of mES culture media. A DNA mixture comprising 9.5 µg of each plasmid—pSUT-REX1, pX335-cas9D10A-Rex1guide4, pX335-cas9D10A-Rex1guide53, and an EF1-GFP construct were prepared in 500uL of OPTI-MEM (Thermo Fisher Scientific, #31985062) with 20 µL P3000 reagent (Thermo Fisher Scientific, #L3000008). In parallel, 20 µL Lipofectamine-3000 (Thermo Fisher Scientific, #L3000008) was diluted in 500 µL OPTI-MEM. The two solutions were combined and incubated for 15 minutes at room temperature before adding to cells. Six hours post-transfection, after confirming cell attachment, half of the medium was replaced with fresh mES cell culture medium to dilute the Lipofectamine-3000. 72h after transfection, GFP-positive single cells (indicating successful transfection) were sorted by FACS and plated in 0.1% gelatin-coated 96-well plates with 200 µL of mES cell culture medium per well. Monoclonal lines were established, expanded and validated by PCR amplification, mCherry imaging (positive signal) and GFP (negative signal) (**Figure S6B**). Homozygous integration at the Rex1 locus was verified, and the resulting cell line was designated RM-CGR8. Potential off-target effects were evaluated by PCR amplification and Sanger sequencing of the top six predicted off-target sites, which were identified using CRISPOR ^38^, confirming that no mutations were introduced.

The YPet-Nanog cell line was generated using CRISPR/Cas9-mediated homology-directed DNA repair (**Figure S6C**). A guide RNA (gRNA) targeting the NANOG N-terminal sequences were designed using CRISPOR ^38^. This gRNA was cloned into the pX330-gRNA vector (a gift from Charles P. Lai; Addgene plasmid #158973) and by digestion with BbsI to construct the NANOG gRNA302. The pSUT45-Nter-cassette-Bsd-P2A-3HA-Ypet-linker was constructed by cloning a PCR-amplified YPet sequence from the pSUT2-TFcassette-puro plasmid (ordered from GeneArt) into the backbone of the pNter-Bsd-mCh-AID plasmid. Briefly, the YPet sequence was amplified using primers with XbaI and NdeI restriction sites. The PCR product and the backbone plasmid were then digested with XbaI (NEB, #R0145S) and NdeI (NEB, #R0111S) restriction enzymes following New England Biolabs’ recommended protocols.

The backbone was dephosphorylated using Antarctic Phosphatase (NEB, #M0289S) following the New England Biolabs protocol. Products were then ligated at a 3:1 insert-to-vector ratio (vector: 3732 bp; insert: 744 bp) using T4 DNA ligase (NEB, #M0202S). The ligation product was transformed into homemade HB-101 competent bacteria. Clones were screened by PCR to confirm the presence of the full YPet sequence, and sequencing was performed to ensure the cassette contained no point mutations. The final plasmid was designated as pSUT45.

Prior to transfection, 80 bp homology arms corresponding to the NH2-terminal part of NANOG were added to the cassette via PCR amplification using the primers Nanog_homoarm_Bsd_for (5’-TTT GGT TGT TGC CTA AAA CCT TTT CAG AAA TCT CTT CTC TCG CCA TCA CAC TGA CAT GGC CAA GCC TTT GTC TCA AG-3’) and Nanog_homarm_endYPetlinker_rev (5’-AAT TCG ATG CTT CCT CAG AAC TAG GCA AAC TGT GGG GAC CAG GAA GAC CCA CAC TCG CCT CCG CCT CCG CCT CCT TAT AG-3’). For transfection, approximately 1.5 million cells were seeded in a 10 cm dish with a total volume of 7 mL of medium. The DNA mixture (ADN MIX Bsd) was prepared by combining 500 µL of OPTIMEM (Thermo Fisher Scientific, #31985062), 0.5 µg of pSUT34 guide RNA302 plasmid, 18.2 µg of the pSUT45-Nter-Bsd-P2A-3HA-Ypet-linker cassette, and 20 µL of P3000 reagent (Thermo Fisher Scientific, #L3000008). Separately, the Lipo MIX was prepared with 500 µL of OPTIMEM (Thermo Fisher Scientific, #31985062) and 20 µL of Lipofectamine-3000 (Thermo Fisher Scientific, #L3000008). The ADN MIX and Lipo MIX were combined and incubated for 15 minutes at room temperature before being added to the cells. Transfected cells were incubated at 37°C, and the transfection efficiency was optimized by using lower volumes. Clones were screened by PCR for insertion of the YPet sequence, and Western Blotting using an anti-Nanog antibody was performed on a selected homozygous knock-in clone to confirm that both alleles were fused to YPet (**Figure S6D**).

### Western Blotting

YPet-Nanog homozygous knock-in cells were lysed in Pierce RIPA Buffer (Cat. 89900, ThermoScientific) supplemented with protease inhibitors and PMSF. The control RM-CGR8 cell line was treated the same way. Samples were mixed with 3X SDS Buffer (Cat. B7703S, NEB) complemented with 10% beta-mercaptoethanol and incubated at 95°C for 5 min. Samples were loaded on a 4-20% Tris-glycine gel (Cat. 4561094, Bio-Rad) together with a PageRuler Plus protein ladder (Cat. 26619, ThermoFisher). After gel separation, proteins were transferred to an EtOH-activated PVDF membrane using the Bio-Rad Trans-Blot® wet transfer system. The membrane was blocked with 5% BSA in PBS with 0.05% Tween-20 (BSA/PBS-T) for 30 minutes at RT, followed by incubation with Rabbit anti-Nanog antibody (Cat. 8822; Cell Signaling) in BSA/PBS-T at 1:1000 overnight at 4°C. The next day, the membrane was washed once with PBS-T and then incubated with anti-Rabbit HRP Conjugate (Cat. W4018, Promega) at 1:10’000 for 45 minutes at room temperature. The membrane was washed four times in PBS-T, developed with Clarity Western ECL Substrate (BioRad), and detected on an Amersham Imager 680 system (GE Healthcare).

### Cell sorting based on TF concentration

For sorting cells with varying TF concentrations, 80 million CGR8 wild-type cells were collected and fixed by incubating them in 2 mM disuccinimidyl glutarate (DSG; BioConcept AG, #3-05F29-I) in PBS (Gibco, #14040091) for 40 minutes at room temperature, followed by incubation in 1% formaldehyde (Thermo Fisher Scientific, #28906) in PBS for 10 minutes at room temperature. Cells were then centrifuged at 500 x g for 5 minutes at 4°C, and washed twice with ice-cold PBS (Gibco, #14040091). Cells were resuspended in ice-cold PBS (Gibco, #14040091) and stained with 50 nM Hoechst 33342 (Thermo Fisher Scientific, #H3570) for 15 minutes while rotating.

From this point, all steps were carried out on ice. Cells were permeabilized by adding ice-cold 70% ethanol in PBS (BioConcept AG, #3-05F29-I) and incubated for 15 minutes. After permeabilization, cells were centrifuged at 500 x g for 5 minutes at 4°C and washed three times with ice-cold PBS (Gibco, #14040091). Cells were then incubated with primary antibodies diluted in ice-cold PBS (Gibco, #14040091) containing 10% ES-cell qualified fetal bovine serum (Gibco, #16141-079). The primary antibodies used were anti-NANOG (1:200, Cell Signaling Technology, #8822S), anti-OCT4 (1:100, Cell Signaling Technology, #5677S), and anti-SOX2 (1:100, Cell Signaling Technology, #23064S). Incubation was conducted for 1 hour at 4°C on an elliptical rotator. After primary antibody incubation, cells were centrifuged at 500 x g for 5 minutes at 4°C and washed twice with PBS (Gibco, #14040091). Cells were then incubated with Alexa-Fluor 647 anti-rabbit (H+L) secondary antibody (Thermo Fisher Scientific, #A-21443) diluted in PBS (Gibco, #14040091) with 10% fetal bovine serum (Gibco, #16141-079) for 40 minutes at 4°C on elliptical rotation. Following secondary antibody incubation, cells were centrifuged at 500 x g for 5 minutes at 4°C and washed twice with PBS (Gibco, #14040091).

For sorting Ypet-NANOG cells at varying NANOG concentration, cells were collected, fixed with DSG and 2% Formaldehyde and stained with 50nM Hoechst 33342 as previously described. The cells were then washed twice with cold-PBS.

Fluorescence-activated cell sorting (FACS) was conducted on a FACSAria II or FACSAria Fusion (BD Biosciences). Cells were transferred to FBS-coated 5 mL polystyrene round-bottom tubes (Falcon, #352235) before sorting. Sorted cells were then collected in FBS-coated collection tubes (Falcon, #352001) containing 250 µL of PBS with 10% fetal bovine serum (Gibco, #16141-079) to prevent adherence. Approximately 2 million cells were sorted per concentration gate. Sorted cells were pelleted by centrifugation at 500 x g for 5 minutes at 4°C and kept at −80°C for chromatin extraction.

### Quantification of OCT4, SOX2, and NANOG in mES Cells by Antibody Staining

Approximately 4×10^7^ CGR8 wild type mES cells were collected to assess OCT4, SOX2, and NANOG concentrations. Cells were fixed by incubation in 2 mM disuccinimidyl glutarate (DSG; BioConcept AG, #3-05F29-I) in PBS (Gibco, #14040091) for 40 minutes at RT, followed by a 10-minute incubation in 1% formaldehyde (Thermo Fisher Scientific, #28906) in PBS at room temperature. Cells were centrifuged at 500 x g for 5 minutes at 4°C and washed twice with ice-cold PBS (Gibco, #14040091). A subset of cells was set aside as a negative control (unstained, fixed cells). The remaining cells were divided into two groups: 2/3 for antibody staining (Cells A) and 1/3 for secondary antibody-only controls (Cells B).

Cells A were stained with 50 nM Hoechst 33342 (Thermo Fisher Scientific, #H3570) for 15 minutes with rotation. Following Hoechst staining, Cells A, Cells B, and negative control cells were permeabilized by adding ice-cold 70% ethanol in PBS (BioConcept AG, #3-05F29-I) and incubating for 15 minutes. After permeabilization, cells were centrifuged at 500 x g for 5 minutes at 4°C and washed three times with ice-cold PBS. All cells were then resuspended in FACS buffer (2% FBS in PBS; Gibco, #10437028) in preparation for antibody staining. Some Cells A were set aside as a Hoechst-only control.

Cells A were incubated with primary antibodies: anti-OCT4 at 5 µg/mL (diluted 1:100; eBioscience, #EM29), anti-SOX2 at 1 µg/mL (diluted 1:200; Santa Cruz Biotechnology, #sc-365823), and anti-NANOG (diluted 1:200, Cell Signaling Technology, #8822S) in FACS buffer for 1 hour at 4°C with elliptical rotation. Following incubation, Cells A were washed three times with ice-cold PBS. Cells B, designated for secondary antibody controls, were similarly resuspended in FACS buffer but were not incubated with primary antibodies.

After washing, both Cells A and Cells B were incubated with secondary antibodies for 45 minutes at 4°C in the dark with elliptical rotation. Cells A were stained with Alexa-Fluor 647 anti-rabbit (20 µg/mL, diluted 1:100; Thermo Fisher Scientific, #A-21443) for NANOG, Alexa-Fluor Plus 555 anti-mouse (20 µg/mL, diluted 1:100; Thermo Fisher Scientific, #A-32727) for SOX2, and Alexa-Fluor 488 anti-rat (20 µg/mL, diluted 1:100; Thermo Fisher Scientific, #A-21208) for OCT4. Cells B were split into three samples, with one of the secondary antibodies added to each sample as a control. After staining, cells were centrifuged at 500 x g for 5 minutes at 4°C and washed three times with ice-cold PBS to remove excess antibodies.

After the final wash, all cells were resuspended in 500 µL of FACS buffer (2% FBS in PBS; Gibco, #10437028) and transferred to FBS-coated 5 mL polystyrene round-bottom tubes (Falcon, #352235) to ensure a single-cell suspension. Compensation controls—including Hoechst-only stained cells, unstained samples, and secondary antibody-only controls for each transcription factor—were prepared in parallel to adjust for autofluorescence and background signal.

Flow cytometry was conducted to quantify the concentrations of OCT4, SOX2, and NANOG across the cell population on a LSRII Analyzer (BD Biosciences). Fluorescence intensity data were analyzed using FlowJo software (BD, #653061) to create the compensation matrix and determine TF concentrations within the mES cell population. This experiment was performed as one replicate.

### Chromatin Immunoprecipitation (ChIP-Seq)

Cells were resuspended in 1 mL of Lysis Buffer 1 (LB1) containing the following components: 50 mM HEPES-KOH, pH 7.4; 140 mM NaCl; 1 mM EDTA (ITW Reagents, #A3145.0500); 0.5 mM EGTA (AG Scientific, #AGS-E-2491-50ML); 10% glycerol (Chemie Brunschwig AG, # FSHG/0650/08); 0.5% NP-40 (Thermo Scientific, #85124); and 0.25% Triton X-100. This buffer was supplemented with a 1:100 dilution of Protease Inhibitor Cocktail (Sigma, #P8340-1ML). Cells were incubated in this buffer for 10 minutes at 4°C with gentle agitation (100 rpm) to ensure efficient lysis. Following incubation, the cells were centrifuged at 1,700 x g for 5 minutes at 4°C to pellet the nuclei. The supernatant was carefully removed to avoid disturbing the pellet. The resulting pellet was then resuspended in 1 mL of Lysis Buffer 2 (LB2), which consisted of 10 mM Tris-HCl, pH 8.0 (ITW Reagents, #A4577); 200 mM NaCl; 1 mM EDTA (ITW Reagents, #A3145.0500); and 0.5 mM EGTA (AG Scientific, #AGS-E-2491-50ML). This buffer was also supplemented with a 1:100 dilution of Protease Inhibitor Cocktail (Sigma, #P8340-1ML). The cells were incubated in LB2 for another 10 minutes at 4°C with gentle agitation (100 rpm) to further facilitate the removal of cytoplasmic components. Following incubation, the nuclei were centrifuged at 1,700 x g for 5 minutes at 4°C, and the supernatant was carefully removed.

To prepare the chromatin, the pellet was washed twice in 0.5 mL of SDS Shearing Buffer, composed of 10 mM Tris-HCl, pH 8.0 (ITW Reagents, #A4577); 1 mM EDTA (ITW Reagents, #A3145.0500); and 0.15% SDS. This buffer was also supplemented with a 1:100 dilution of Protease Inhibitor Cocktail (Sigma, #P8340-1ML). After each wash, the cells were centrifuged at 1,700 x g for 5 minutes at 4°C, with the cell pellets left undisturbed during each wash (i.e., they were not resuspended between washes). Finally, the pellet was resuspended in 450 µL of fresh SDS Shearing Buffer. Chromatin was then sonicated using a Covaris E220 ultrasonicator with the following settings: 20 minutes at a 5% duty cycle, 140 W, and 200 cycles. This shearing process fragmented the chromatin to sizes suitable for further downstream applications. After sonication, the chromatin solution was centrifuged at 10,000 x g for 5 minutes at 4°C to remove any debris, and the supernatant containing the sheared chromatin was transferred to new 1.5 mL Eppendorf tubes (Eppendorf, #0030108051).

ChIP was performed using the ChIP-IT High Sensitivity Kit (Active Motif, #53040) according to the manufacturer’s protocol. For each TF ChIP-seq sample, 10 ng of Drosophila spike-in chromatin (Active Motif, #53083) and 0.5 µg of spike-in antibody (anti-H2Av, Active Motif, #61686) were added to 3.5 µg of chromatin to ensure consistency and normalization between samples. The following primary antibodies for TF-targeted ChIP-seq were used: 3.1 µL of OCT4 antibody (Cell Signaling Technology, #5677S), 3.1 µL of SOX2 antibody (Cell Signaling Technology, #23064), or 1.55 µL of NANOG antibody (Cell Signaling Technology, #8822). ChIP-seq libraries were prepared using the NEBNext Ultra II DNA Library Prep Kit (New England Biolabs, #E7645) following the manufacturer’s protocol. Libraries were sequenced on an Illumina NextSeq 500 system using paired-end 75-bp reads to obtain high-resolution mapping of TF binding sites across the genome. Two replicates of ChIP-seq experiments were performed on NANOG, OCT4 and SOX2 with intracellular antibody staining. ChIP-seq experiments on YPet-NANOG were performed as one replicate.

### ATAC-seq on fixed cells

ATAC-seq was performed on fixed and sorted wild-type (WT) CGR8 mouse embryonic stem (mES) cells. Cells were prepared in the same manner as for ChIP-Seq, including fixation with 2 mM DSG (Disuccinimidyl glutarate, Thermo Fisher Scientific, #20593) for 40 minutes, followed by a 10-minute incubation with 2% formaldehyde (Thermo Fisher Scientific, #28908) to ensure cross-linking of chromatin. After fixation, cells were stained with Hoechst 33342 (Thermo Fisher Scientific, #H3570) for DNA content and immunostained for SOX2 or OCT4 to perform cell sorting recreating the same gating strategy than for ChIP-sequenced sorted cells. The Fixed Cell ATAC-Seq Kit (Active Motif, #53151) was used following the manufacturer’s protocol. For each ATAC-Seq reaction, 100,000 cells were used. ATAC-seq experiments were performed as one replicate.

### ChIP-seq and ATAC-seq analysis

Sequencing results from the newly generated ChIP-seq libraries were aligned to the Mus musculus mm10 genome (GRCm38, release M25) and the Drosophila melanogaster genome (BDGP6, release 28) using STAR version 2.7.0e ^39^ with parameters optimized for alignment: --alignMatesGapMax 2000 --alignIntronMax 1 --alignEndsType EndtoEnd -- outFilterMultimapNmax 1. Duplicate reads, as well as reads that did not map to chromosomes 1-19 or X and Y, were removed using Picard (Broad Institute), and all processed BAM files were indexed using SAMTools (v1.10) ^40^.

For peak calling, MACS2 (v2.2.4) ^41^ was employed with the parameters -f BAMPE -g mm, specifically for the mouse genome, while peaks within ENCODE blacklist regions for mm10 ^42^ were excluded to prevent artifacts. BigWig files were created from the down-sampled BAM files using the bamCoverage function from deepTools (v3.5.1) ^43^ with the –normalizeUsing RPKM parameter to ensure consistent RPKM-normalized enrichment scores.

To perform spike-in normalization for ChIP-seq, Drosophila read counts were quantified within each BAM file, and normalization factors were calculated to equalize read counts across all Drosophila BAM files. These normalization factors were then applied to down-sample the mouse genome-aligned BAM files using SAMTools, ensuring consistent read depth across samples. For comparative analysis across conditions, a comprehensive BED file consolidating all peaks detected in at least one condition was generated as a consensus genome, enabling direct comparison across samples. Enrichment values within these regions were calculated using the multiBigwigSummary function in deepTools. ATAC-seq data were processed similarly, with down-sampling based on the total number of reads to ensure consistent normalization across samples.

For samples with two replicates, each set of four concentration samples per replicate was normalized together. The normalized BED files per concentration condition were then merged to retain all peaks detected in at least one replicate. BigWig files from both replicates for each condition were averaged using the BigWig merge function in deepTools to create combined profiles for analysis.

### Unsupervised clustering

Binding scores of OCT4 and SOX2 across their target regions were calculated for each concentration condition. For each region, scores were Z-scored across concentrations to place profiles on a common scale and minimize amplitude-driven bias. The optimal number of clusters was estimated in R with gap statistics (factoextra::fviz_nbclust, k-means) (https://rpkgs.datanovia.com/factoextra/index.html, https://cran.r-project.org/web/packages/factoextra/index.html), evaluating k from 1 to 10. The analysis was unsupervised because no labels or prior groupings were provided; clusters were inferred solely from the structure of the normalized data. Once the optimal number of clusters was obtained, clusters were visualized with the pheatmap R package (https://CRAN.R-project.org/package=pheatmap) using the fixed k cluster number. The gap-statistic approach indicated k = 3 for OCT4 and k = 5 for SOX2. The five-cluster solution for SOX2 added little interpretative value, so k = 3 was chosen for SOX2 based on inspection of the gap curve, which provided a clearer summary of SOX2 binding patterns.

### Motif and Sequence enrichment analysis

Motif enrichment was performed using the SEA (Sequence Enrichment Analysis) (Bailey bioRxiv 2021) algorithm from the MEME suite with the JASPAR non-redundant core database 2020 as the reference with addition of NANOG UN0383.1 motif. Default parameters were applied to identify enriched motifs in the ChIP-seq data. Additionally, motif occurrence analysis was conducted using FIMO (Find Individual Motif Occurrences) with a p-value threshold of 0.01 to locate specific motif sites within target regions. For this analysis, FIMO was employed to identify precise binding sites of transcription factors using reference motifs for ESRRB (MA0141.1), NANOG (UN0383.1), NR5A2 (MA0505.1), OCT4 (MA1115.1), the OCT4-SOX2 heterodimer (MA0142.1), and SOX2 (MA0143.1). FIMO scans each sequence in the input file, calculating the likelihood of observing each motif at specific positions, enabling high-resolution identification of binding sites within the ChIP-seq peaks. This approach allows to determine the likelihood of having a motif corresponding to consensus sequence in the peak sequence. FIMO and SEA results were processed using custom R (R Core Team. R: A language and environment for statistical computing. (2020) scripts to filter, organize, and visualize motif enrichment data. Key R packages included tidyverse (Wickham J Open Source Software 2019) for data handling, dplyr (Wickham 2023) for filtering, ggplot2 (Wickham 2009) for visualization, and plyr (Wickham 2011) for data aggregation. SEA results were grouped by experimental condition to identify shared motifs across TFs and conditions, while FIMO data was further refined to highlight significant motifs across clusters. Enrichment scores and motif occupancy were visualized in summary plots, supporting comparative analyses of transcription factor binding across regions.

### Peak annotation and Gene Ontology enrichment analysis

Annotation of peaks to corresponding genes was performed using ClusterProfiler (https://doi.org/10.1089/omi.2011.0118), assigning the nearest gene to each peak. The obtained list of genes was then analyzed for Gene Ontology enrichment using Homer ^44^ and results corresponding to biological process stem cell maintenance and WikiPathway PluriNetwork were selected. To determine enrichment for genes associated with pluripotency maintenance we used a custom list of 24 genes presented in Figure S5.

### TOBIAS analysis

For TOBIAS analysis (Bentsen Nature Comm 2020), ATAC-seq called peaks were overlapped with the SOX2 or OCT4 consensus set of regions derived from the ChIP-seq data to select for accessible regions bound respectively by SOX2 or OCT4. To account for Tn5 transposase insertion bias, bias correction was performed using TOBIAS ATACCorrect on the overlapping files, while excluding regions blacklisted by ENCODE for the mm10 genome. Footprint scoring was then conducted within these regions using TOBIAS FootprintScore to identify protected sites. Finally, TOBIAS BINDetect was applied to compare motif activity across conditions at increasing SOX2 or OCT4 concentrations, using the non-redundant core motif list from JASPAR2022, with the NANOG motif (UN0383.1) manually added to the list for comprehensive analysis.

## Acknowledgments

We thank the Gene Expression, Flow Cytometry, and BioImaging core facilities at EPFL, and Dr Ludovica Vanzan for providing the PE score calculation script, assistance with data analysis, and critical reading of the manuscript. We are thankful for funding by the Swiss National Science Foundation grant CRSII5_189910 and #310030_212197 to DMS.

## Authors Contributions

Conceptualization, RM and DMS; Methodology, RM, AT and DMS; Formal Analysis, RM, AT, and DMS; Investigation, RM, AT and CD; Resources, DMS. Funding Acquisition, DMS. Supervision, DMS.

## Supplementary figures

**Supplementary Figure 1.**
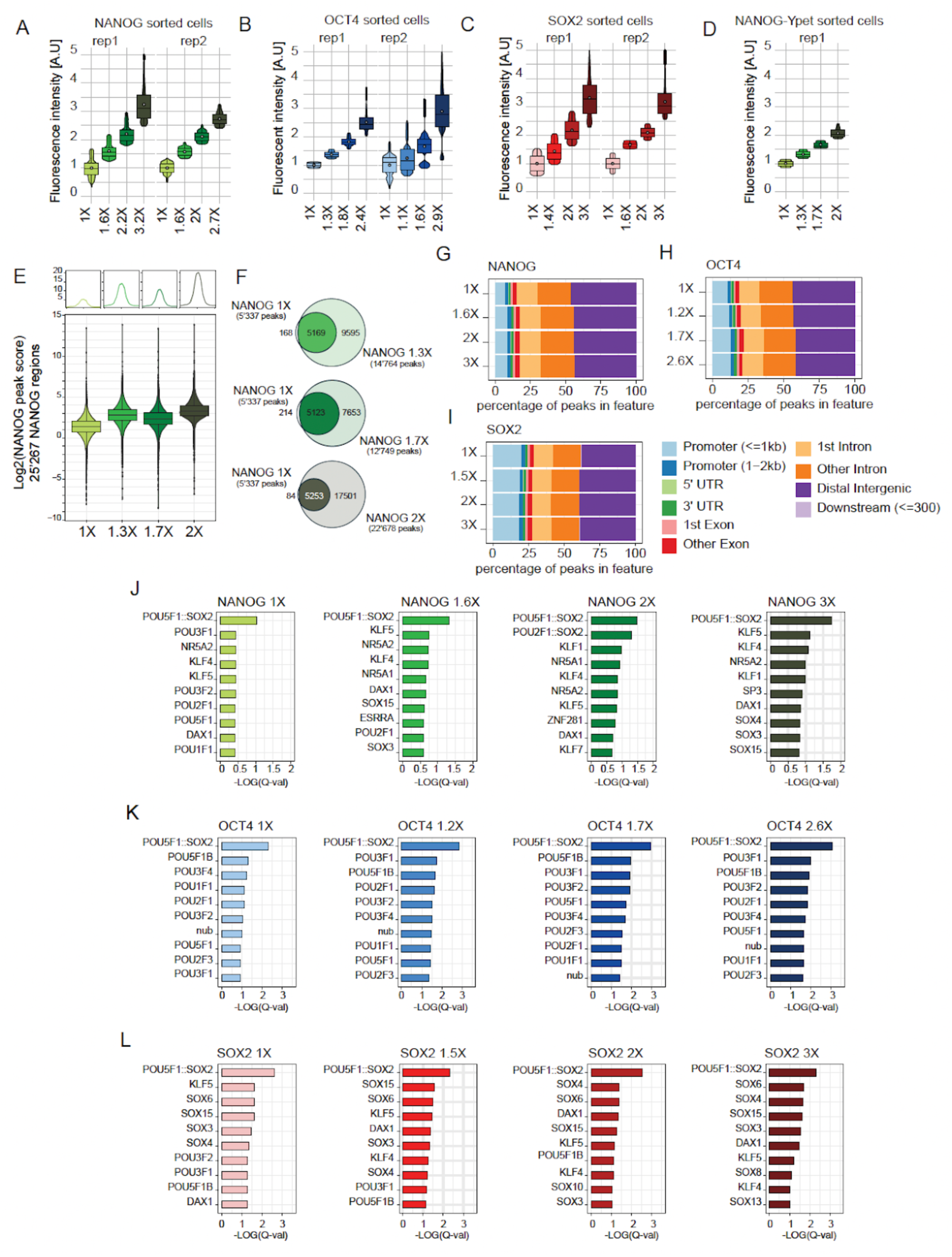
(A-C) Fluorescent intensities of individual replicates of the sorted cell populations for NANOG (A), OCT4 (B), and SOX2 (C). (D) Distribution of fluorescence intensity in the sorted populations YPET-NANOG. The fold change is calculated as explained in Figure 1. The white dot represents the average intensity per gate. (E) Representation of YPET-NANOG binding site occupancy as its concentration increases. Boxplots as in Fig 1. (F) Overlap of bound regions at different concentrations for YPET-NANOG. (G-I) Distribution of bound sites across different types of regions for NANOG (G), OCT4 (H) and SOX2 (I). (J-L) List of the top 10-enriched motifs enriched in regions bound by (J) NANOG, (K) OCT4, or (L) SOX2 as their concentrations increase.

**Supplementary Figure 2.**
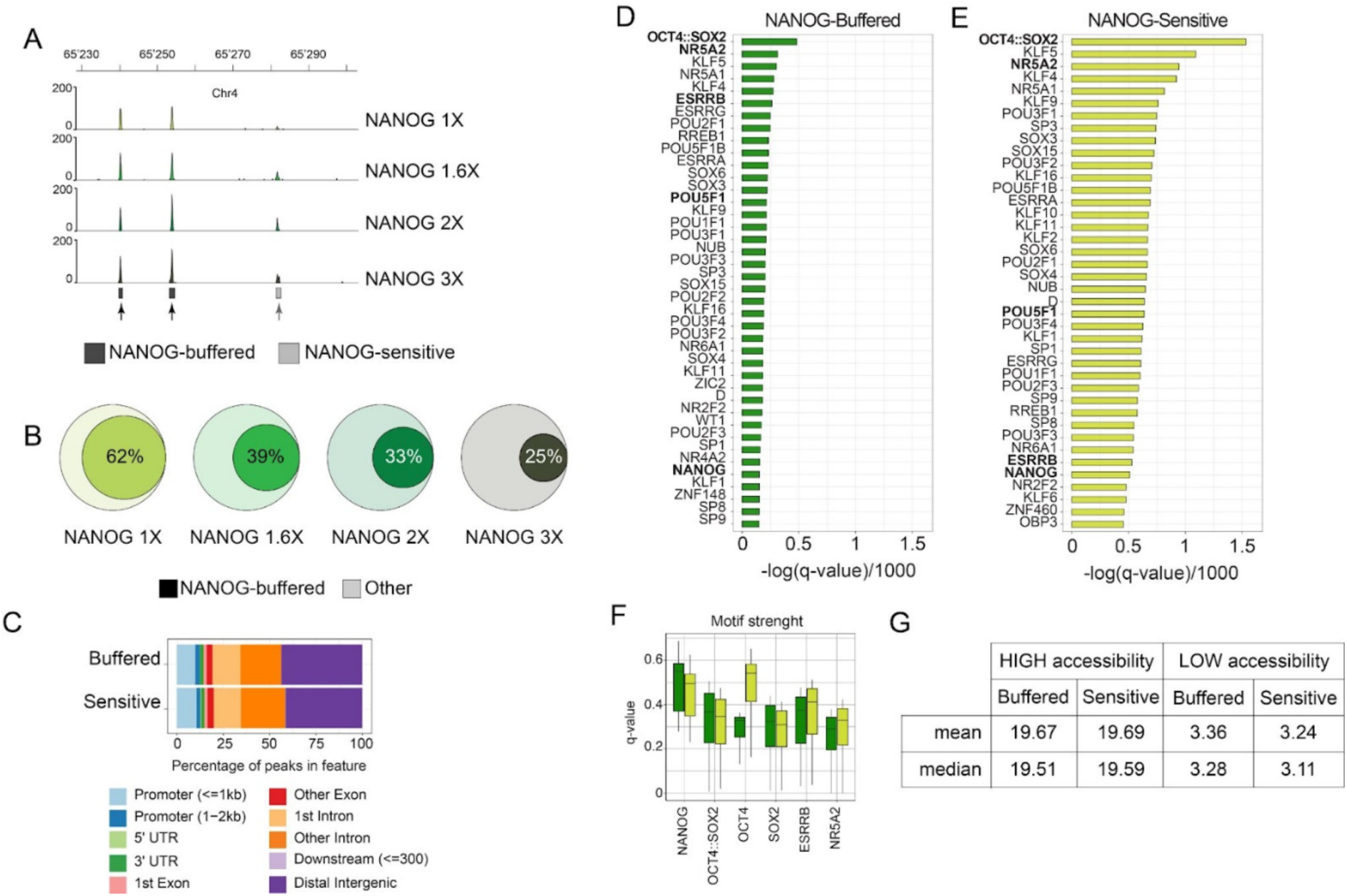
(A) Genome tracks of NANOG ChIP-seq on the sample with different NANOG concentrations, showing examples of buffered and sensitive regions. (B) Overlap of buffered NANOG regions with all bound regions at different NANOG concentrations. (C) Distribution of bound sites across different types of regions for NANOG-buffered and NANOG-sensitive regions. (D-E) Top 40 TF motifs enriched in NANOG-buffered (D) and NANOG-sensitive (E) regions. (F) Motif quality of NANOG, OCT4, SOX2, ESRRB and NR5A2 in NANOG-buffered and NANOG-sensitive regions. (G) Average and mean of the accessibility scores of the different regions selected for analysis in Figure 3C-D.

**Supplementary Figure 3.**
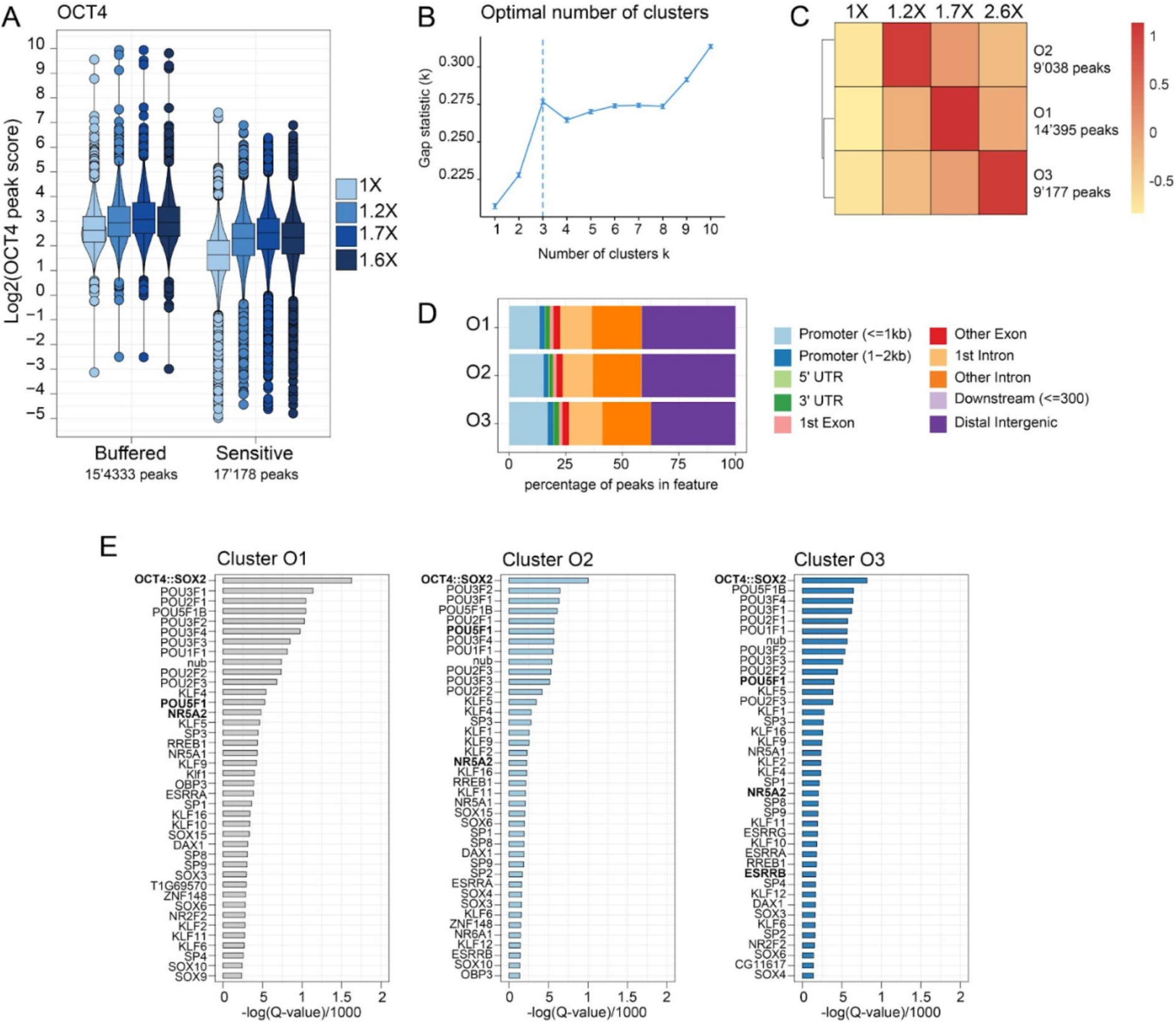
(A) Distribution of OCT4 peak scores at OCT4-buffered and OCT4-sensitive regions at varying concentrations. Boxplots as in Fig 1. (B) Gap statistics for OCT4 clusters. (C) Heatmap of Z-scored signal across three OCT4 peak clusters at OCT4 concentrations (1X, 1.5X, 2X, 3X). Each cell reports the mean Z-score across peaks in the cluster. The color scale ranges from 1 (red) to −0.5 (yellow). Row labels O1–O3 include the number of peaks per cluster. The row dendrogram reflects hierarchical clustering of the clusters by their concentration response. (D) Distribution of bound sites across different types of regions for the three OCT4 clusters. List of the top 40-enriched motifs enriched in regions bound by OCT4 for the three clusters.

**Supplementary Figure 4.**
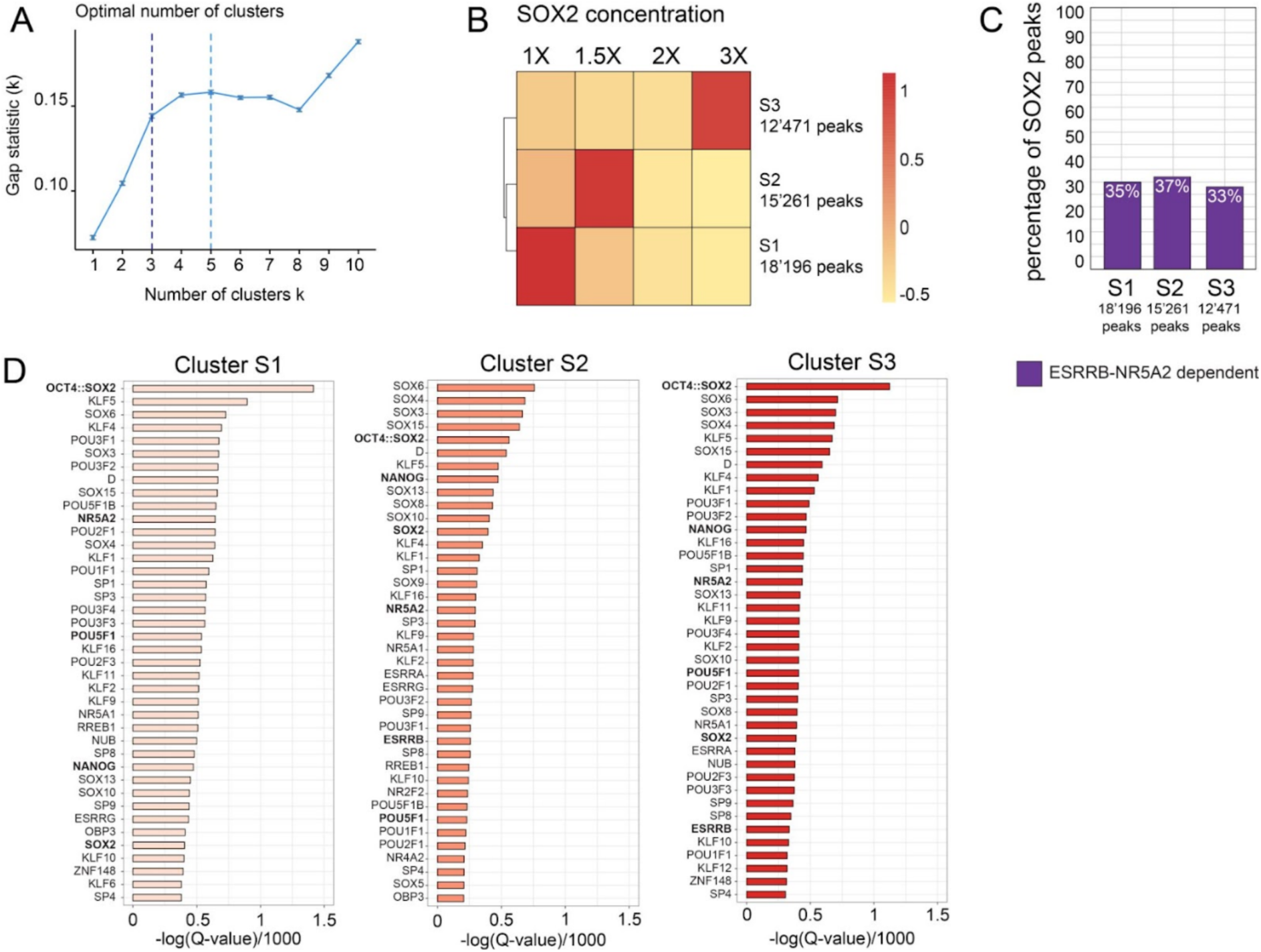
(A) Gap statistics for SOX2 clusters. (B) Heatmap of Z-scored signal across three SOX2 peak clusters at four SOX2 concentrations (1X, 1.5X, 2X, 3X) (as in Figure S3C). (C) Proportions of ESRRB-NR5A2-dependent regions in the three SOX2 clusters. (Festuccia et al., 2021) (D) List of the top 40-enriched motifs enriched in regions bound by SOX2 for the three clusters.

**Supplementary Figure 5.**
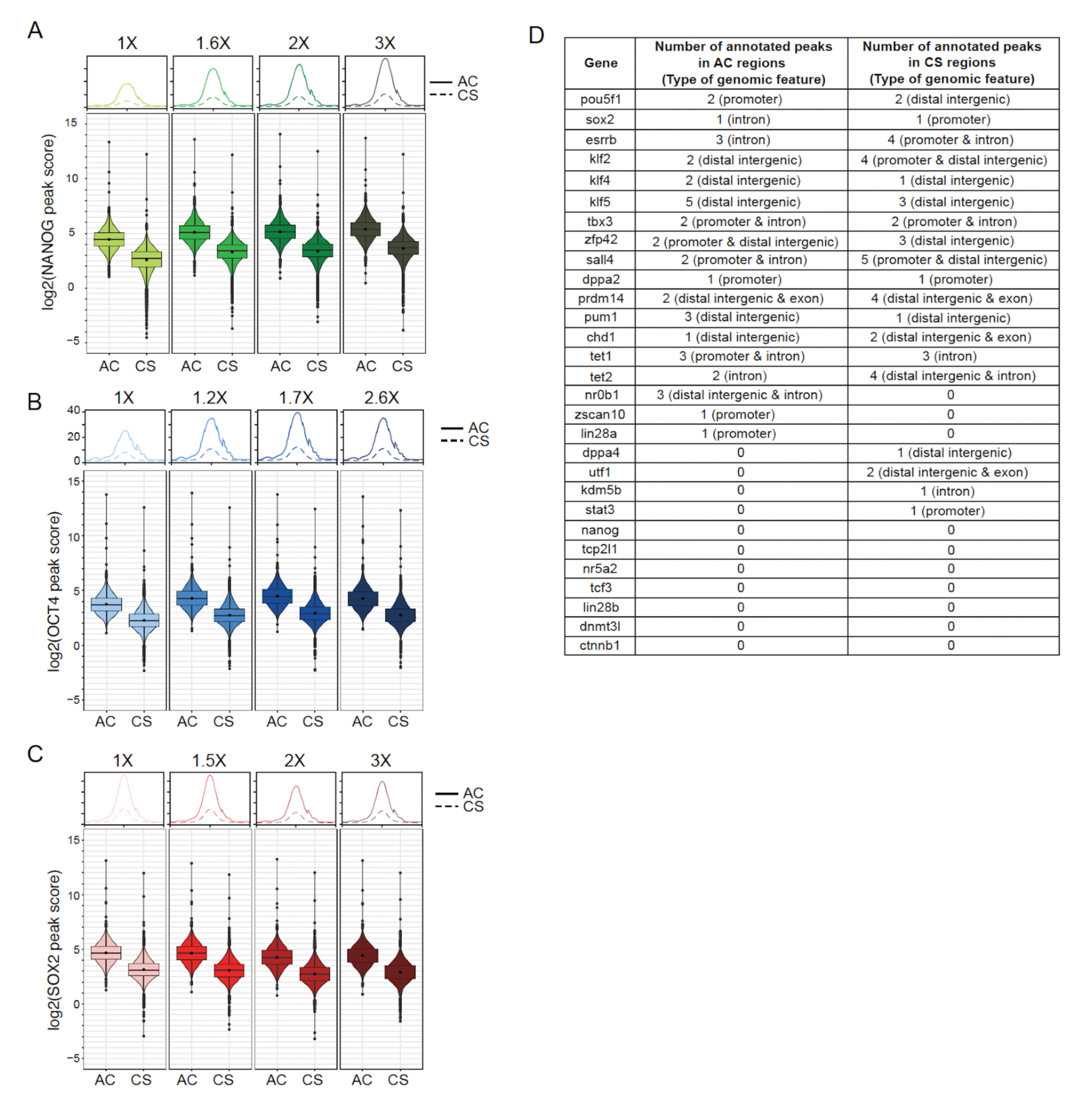
NANOG (A), OCT4 (B) and SOX2 (C enrichment for the four concentration bins, in the AC and CS regions. Boxplot as in Fig 1. D. List of pluripotency genes with AC or CS annotated peaks. Genes were assigned to a given peak by proximity.

**Supplementary Figure 6.**
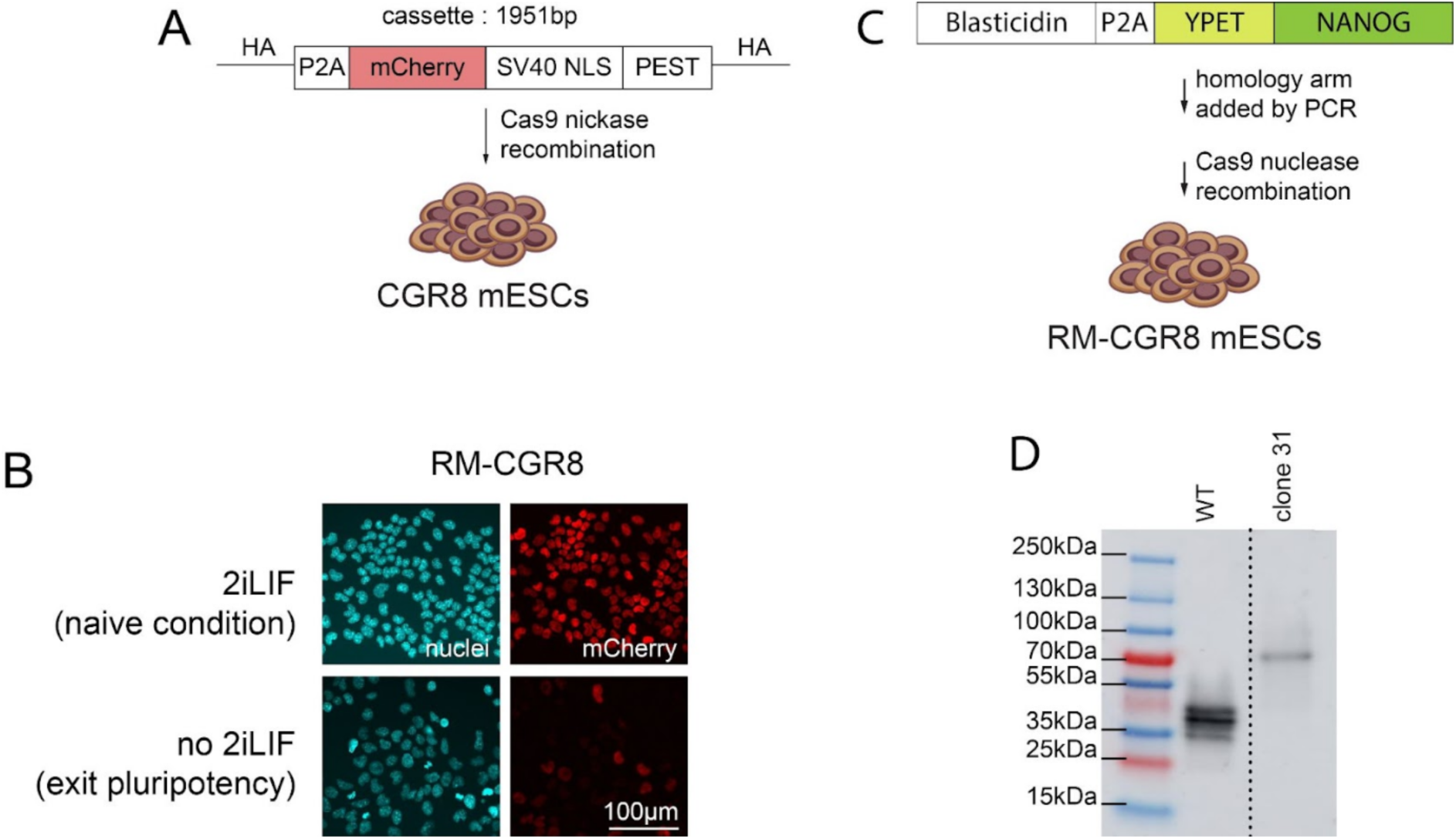
(A) Schematic representation of RM-CGR8 cell line generation. mCherry, was inserted at the C-terminus of naive pluripotency marker REX1 using CRISPR/Cas9 DNA repair strategy. Briefly, CGR8 WT cells were transfected with a P2A-mCherry-NLS-PEST cassette flanked by 500bp homology arms and a pair of CRISPR/ Cas9 nickase to induce double strand break repair and promote none-homologous-end-joining (NHEJ) repair. The cell line was named R(ex1)-M(cherry)-CGR8. (B) Fluorescence imaging of RM-GCR8 cells kept in naive condition (top) expressing REX1 and therefore mCherry positive, and 72h post-removal of 2i+LIF from the culture medium, allowing cells to exit pluripotency (bottom), REX1 and mCherry negative. (C) Schematic representation of the Ypet-NANOG cell line generation. Sequences corresponding to the NANOG C-terminus coding sequence were added by PCR amplification to a blasticidin-P2A-YPET cassette prior to transfection in RM-GCR8 cells with a CRISPR/ Cas9 nuclease. (D) Western blotting using an anti-NANOG antibody, demonstrating that all NANOG proteins are fused to YPet.

